# Suspected distortion of citations in high-impact cancer journals

**DOI:** 10.64898/2026.05.25.727627

**Authors:** Baptiste Scancar, Jennifer A. Byrne, David Causeur, Adrian G. Barnett

## Abstract

Research and scholarship are shaped by article citations, which underpin the communication of ideas, assignment of credit, journal impact factors, and author career progression. Given their key influence on author and journal metrics, citations can be intentionally manipulated to inflate the reputation of journals and researchers. Paper mills, unethical organisations that produce and sell manuscripts and publishing services, may also be manipulating citations, but the extent of this manipulation is unknown. Here, we show that molecular cancer articles sharing features with retracted papers from paper mills display citation patterns that suggest systematic inflation. These articles were published in journals in the top decile of journal rankings. Suspected paper mill articles received 50 to 100% more citations than other papers 1 to 3 years after publication, while paradoxically attracting fewer readers and online accesses. Suspected paper mill articles also cited – and were cited by – other suspected paper mill articles, which frequently appeared in journals previously reported as paper mill targets. Citations from suspected paper mill articles measurably inflated journal citation metrics. These findings suggest that paper mills inflate the citation metrics of supported publications and affected journals at scale. This widespread manipulation of citation metrics may amplify unreliable findings, slowing scientific progress and spurring unreasonable citation benchmarks for research articles, journals and authors. Our findings point to measurable signatures of citation manipulation that could support article- and journal-level screening.

**Significance statement:** The central role of citations in research evaluation makes them an attractive target for gaming. It has been suggested that fraudulent organisations, known as *research paper mills*, may manipulate citations for a price. This research uncovers the impact of potential paper mill manipulation on citation metrics. Across leading cancer journals, clusters of interconnected papers bearing the hallmarks of paper mill products accumulate far more citations than other papers despite having fewer readers. In some cases, this effect is large enough to inflate journal rankings, highlighting the need for safeguards against paper mills. This study provides evidence that citation manipulation leaves measurable traces in citation patterns, offering a practical basis for future screening and intervention.

## Introduction

Academia operates in a highly competitive, tournament-like environment (1) – where rewards include research grants, academic positions and career progression (1, 2). The volume of scientific output has grown exponentially in recent years (3) – at a rate that outpaces economic growth (4) – thereby intensifying the pressure on researchers to stand out. Potentially small differences in commonly used evaluation metrics – such as publication numbers, journal impact factors and citation counts – can translate to advantages, such as securing promotions and funding (1, 5). This reliance on quantitative performance indicators creates strong incentives to maximise article and citation numbers, with unintended consequences including research fraud (6, 7).

*Research paper mills*, organisations providing researchers with fabricated manuscripts for a price (8, 9), are a manifestation of this hyper-competition (8–11). Paper mill manuscripts may include textual and structural similarities (12), report fake data and experiments (9), and contain manipulated or duplicated images (13). These practices are likely to reflect business models reliant on rapid and high-volume manuscript production (8, 14) – which is difficult to reconcile with the slow and costly process of genuine research. In a recent survey of Chinese medical residents, close to half of the respondents admitted to having bought or sold papers or authorships, indicating a widespread acceptance of the transactional nature of authorship (15). Paper mills are growing in both sophistication and scale, with reports describing the involvement of agents and networks that extend beyond manuscript production to influence editorial and publishing processes (16–18). The number of papers retracted due to paper mill concerns (19, 20), as well as the number of papers exhibiting paper mill hallmarks has surged in recent years (21, 22). While paper mills could target low-impact journals and venues with weaker editorial and peer-review controls (11, 16), evidence suggests that high-impact journals are also vulnerable to paper mills (21, 23).

Citations are an important academic currency (24, 25) that is used to evaluate scientists (2, 26), making citations an important target for paper mills and their customers (27). Papers produced by paper mills may cite other paper mill papers (13, 28) and groups of papers that systematically cite one another – termed “*citation cartels*” (29) – have been linked to paper mills (28). Some paper mill publications have attracted considerable citation attention (23, 28, 30) and have been included in systematic reviews (31, 32). At the journal level, citation malpractices associated with paper mills could inflate journal impact factors (27), and concerns about citation and publication practices have contributed to the delisting of journals from Clarivate’s Web of Science (33, 34).

Concerns about manipulated publications are particularly relevant in cancer research, as this field has been identified as being affected by paper mills (12, 21, 35), especially in molecular oncology (8, 12, 36). Suspected paper mill papers have been published in high-impact molecular cancer journals with some accumulating high citation counts (23). As paper mill papers are not genuine scientific contributions, such citation counts may reflect artificial amplification rather than genuine scientific engagement. Artificial citations may distort the evidence base by misinforming future research and clinical translation.

Here we focus on high-impact molecular oncology journals, applying our previous detection model (21) to flag papers with titles and abstracts textually similar to those of retracted paper mill papers, hereafter referred to as “*flagged papers*”. We show that flagged papers, which represent a large percentage of publications in some journals, accumulate many more citations than non-flagged papers, yet paradoxically attract fewer readers. Flagged papers predominantly cite and are cited by other flagged papers, where the resulting citations can measurably inflate the citation metrics of cancer research journals.

## Results

### Flagged papers cluster within leading molecular cancer journals

We defined high-impact molecular oncology journals as cancer journals within the top decile of SCImago (37) rankings that publish molecular oncology research, giving 20 journals and 33,159 index papers published between 2012 and 2023 (SI Appendix, Table S1). Journals were ranked using SCImago’s citations per document (2 years) metric, and molecular cancer research papers were identified within each journal using a validated classification model. We structured our analysis around these index papers, distinguishing the papers they reference (referenced papers) and papers that cite them (citing papers) and classifying each group as flagged or non-flagged using our detection model (21) (Fig. 1). The model to flag suspected paper mill papers was trained on titles and abstracts of retracted paper mill papers and produced a probability score indicating textual similarity to known paper mill publications. Predictions for the flagged papers were made using two probability thresholds (0.6 and 0.9) to assess the robustness of observed patterns to a stricter classification of flagged papers, with the higher threshold prioritising specificity and reducing false positives.

**Fig. 1:**
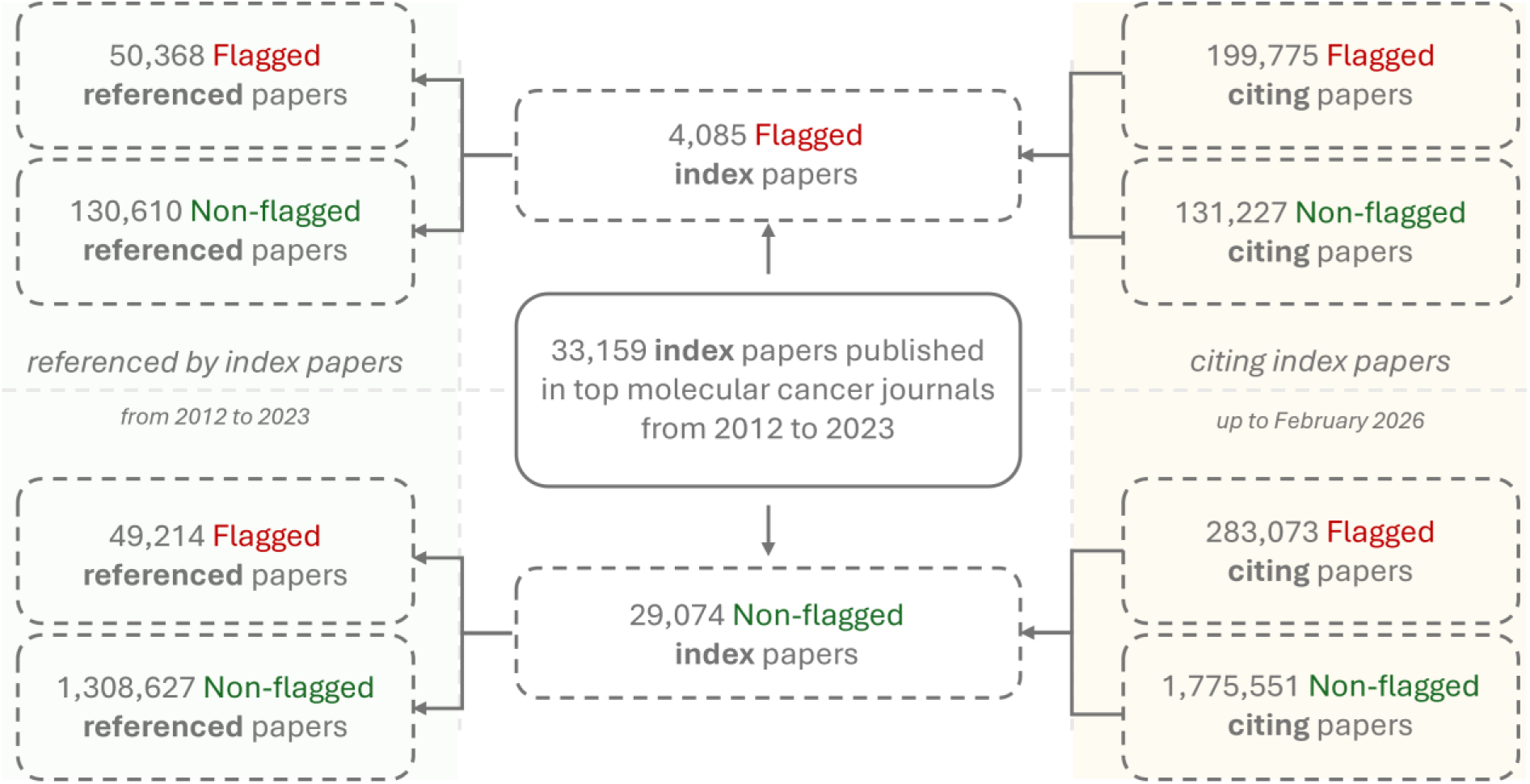
Study design and citation analysis. Index papers (n = 33,159) published between 2012 and 2023 represent the core dataset and were classified as flagged or non-flagged; the 0.6 probability threshold is shown here for illustration. For each index paper, we identified referenced papers (papers cited by index papers, n = ∼1.5 million) and citing papers (papers citing index papers, n = ∼2.4 million), which were also classified as flagged or non-flagged using the same model and 0.6 threshold. Citation flows were then analysed across these groups.

A total of 4,085 index papers (12.3%) were flagged by the model (21) as textually similar to retracted paper mill papers at the 0.6 probability threshold. At the more conservative 0.9 threshold, 933 index papers (2.8%) were flagged. Flagged papers were not evenly distributed across journals but instead clustered in a subset of titles including *Journal of Experimental & Clinical Cancer Research* and *Molecular Cancer*, whereas other journals were only marginally affected (Fig. 2; SI Appendix, Fig. S1, Table S2). Flagged papers were identified across 19 out of 20 journals regardless of their ranking (see rankings in Box 1), with half of the top ten journals having at least 20% of their papers flagged (SI Appendix, Table S2). Authors affiliated with Chinese institutions accounted for 70% of flagged papers, despite representing only 21% of all index papers examined (SI Appendix, Table S3).

**Fig. 2:**
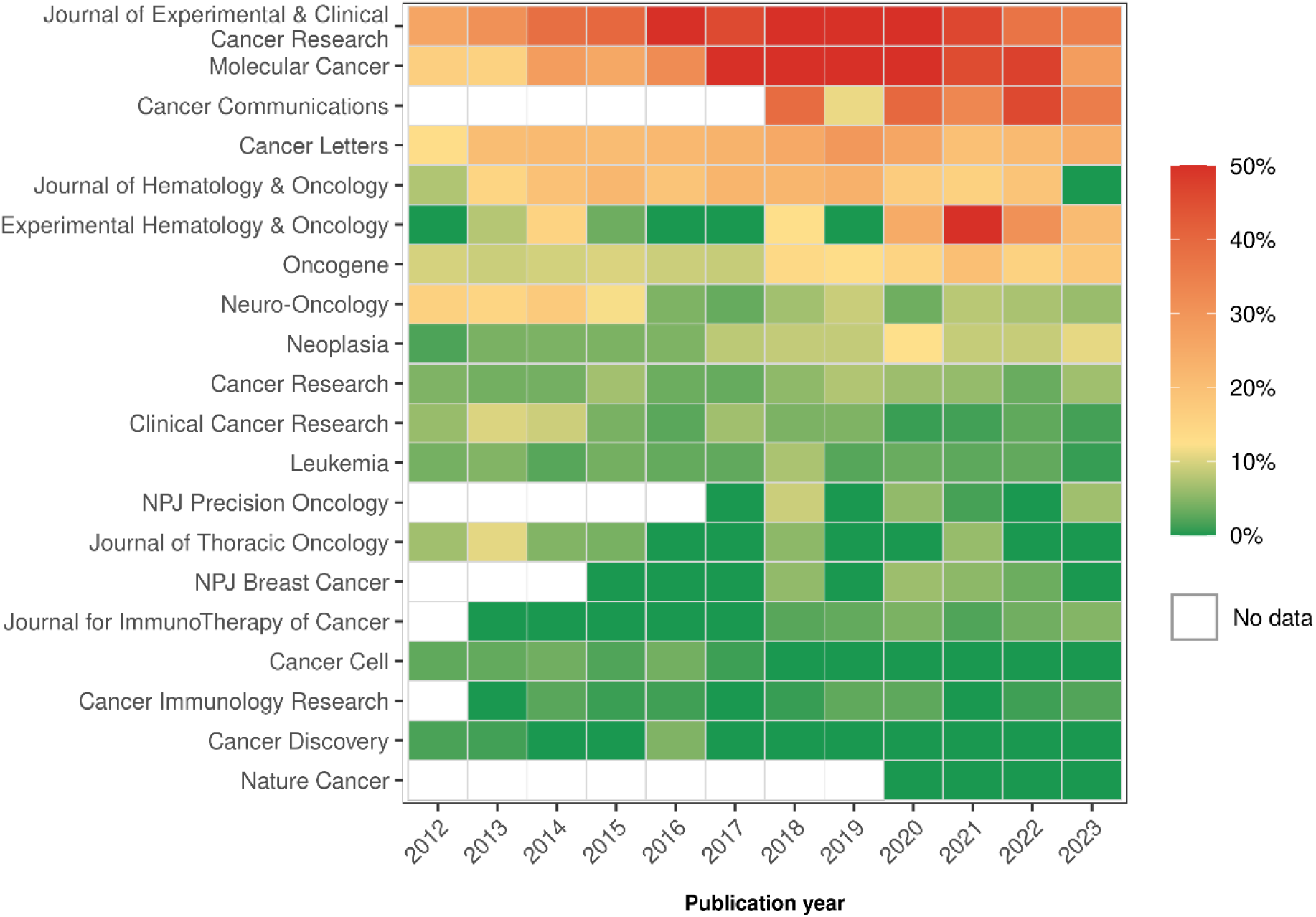
Heatmap showing yearly percentages of flagged papers across leading molecular cancer research journals (0.6 paper mill probability threshold). Journals are ordered according to their overall mean percentage of flagged papers during the study period. The colour scale is centred at 12.3%, which is the overall percentage of flagged papers. Blank cells indicate missing data, either because the journal had not been established or because no molecular cancer research papers were published in that year in that journal. Values are capped at 50%, with all percentages over 50% displayed at the maximum colour intensity. These patterns hold at the conservative 0.9 threshold (SI Appendix, Fig. S1).

Flagged paper percentages of 10 to 20% were observed in some journals in 2012, pointing to an early and sustained presence of flagged papers (Fig. 2). These percentages increased over time in journals with high initial levels, while titles such as *Oncogene*, *Neoplasia*, and *Journal for ImmunoTherapy of Cancer*, which were initially minimally affected, began to show higher percentages from 2018 onwards. A few journals had percentages close to zero in all years. In summary, flagged papers are not randomly distributed but instead cluster within specific high-impact journals.

To provide further evidence that flagged papers are likely paper mill products, we investigated whether they were more likely to be flagged by two external sources of research integrity concerns: PubPeer comments and the Problematic Paper Screener (38) (PPS). We retrieved PubPeer comments for all 4,085 flagged papers at the 0.6 threshold, alongside a comparison subset of 4,088 non-flagged papers sampled proportionally to each journal’s number of non-flagged papers. Flagged papers received PubPeer comments at more than three times the rate of non-flagged papers (18.8% vs. 5.4%) and were nearly four times as likely to receive comments raising direct integrity concerns (17.0% vs. 4.5%). Consistently, 30% of flagged index papers were also flagged by the PPS, compared with only 8% of non-flagged index papers (see SI Appendix for details).

### Flagged papers receive more citations but lower attention

To quantify scientific engagement with index papers, we retrieved citation counts from OpenAlex (39) at three post-publication time points: one year, three years, and since publication. As complementary indicators of attention, we retrieved *Mendeley* readers and the number of online accesses. Using negative binomial regression models, we quantified the rate ratio of these outcomes for flagged versus non-flagged index papers. Given the disproportionate representation of Chinese institutions among flagged papers, results were also assessed after adjustment for China versus non-China authorship to ensure that observed differences were not driven by citation practices specific to authors in China.

Flagged index papers had 50 to 100% higher citation counts than non-flagged papers at both probability thresholds, a finding that remained robust after adjustment for Chinese authorship (Fig. 3; SI Appendix, Fig. S2, Table S4). These findings were also robust when using an alternative citation database (Dimensions.ai) with near-identical rate ratios (SI Appendix, Fig. S3). The citation rate ratio of flagged to non-flagged papers was highest in the early years after publication, reaching an over 100% increase at the 0.9 threshold, before attenuating to an approximately 50% increase over time. In contrast, *Mendeley* readers and online accesses were approximately 10 to 20% lower for flagged than for non-flagged papers, which is the opposite of the expected pattern, as more citations should arise from more readers.

**Fig. 3:**
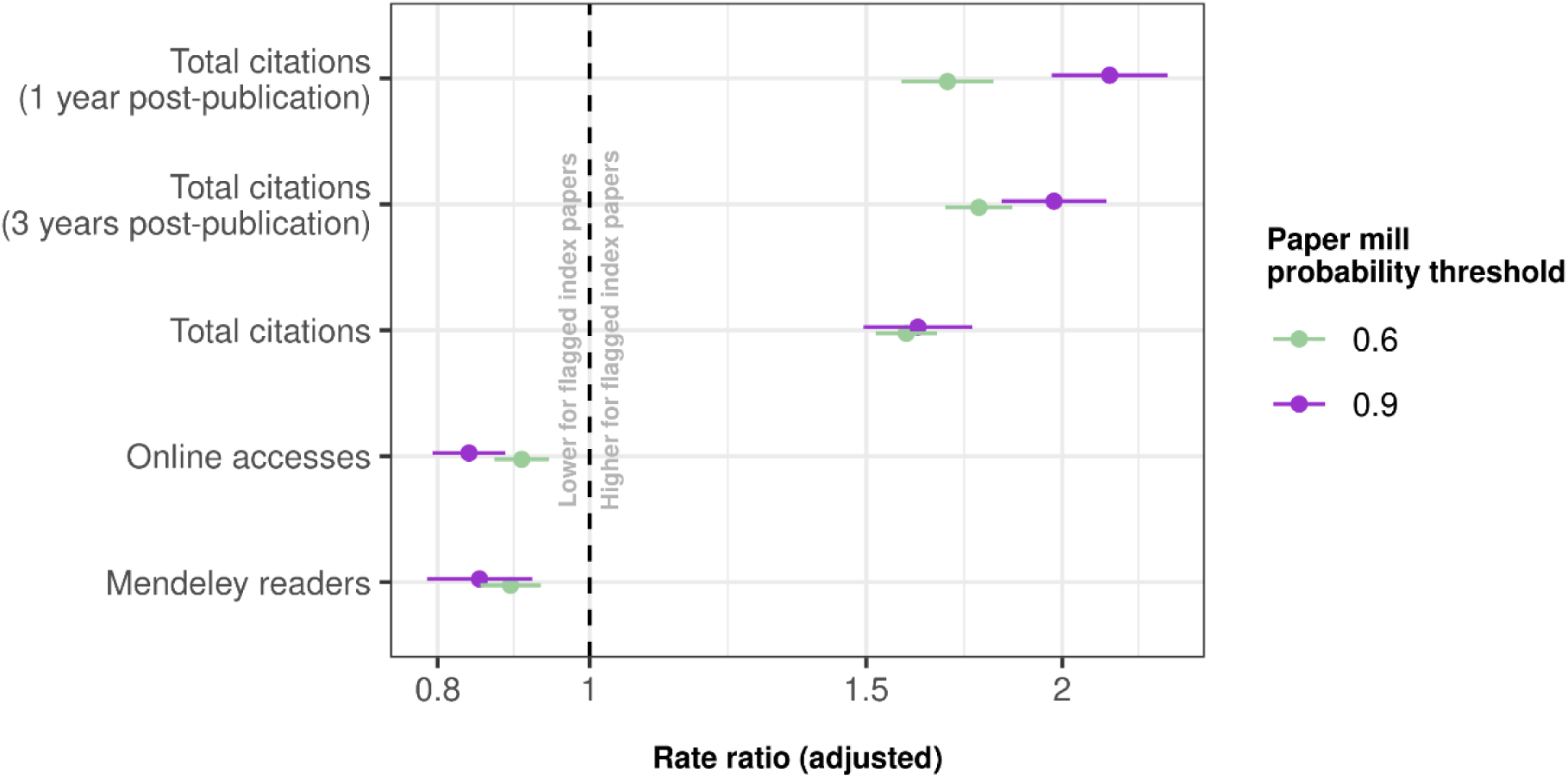
Adjusted rate ratios (RR) for citation and attention outcomes associated with flagged index papers. Attention outcomes (Mendeley readers and online accesses) reflect total cumulative attention since publication. Points represent adjusted rate ratios estimated from negative binomial regression models, comparing flagged and non-flagged index papers based on the paper mill probability threshold (0.6 and 0.9). Models were adjusted for journal and year of publication. Horizontal bars are 95% confidence intervals with cluster-robust standard errors (Journal×Year). Rate ratios are shown on a logarithmic scale. The vertical dashed line indicates a rate ratio of 1 (no difference between flagged and non-flagged papers). The corresponding figure with additional adjustment for Chinese authorship is presented in SI Appendix, Fig. S2. Adjusted rate ratios produced using Dimensions.ai data are presented in SI Appendix, Fig. S3. A sensitivity analysis restricted to papers mentioning non-coding RNA keywords in their titles and / or abstracts is presented in SI Appendix, Fig. S4.

Considering that paper mills are known to target non-coding RNA research (16, 27) – a subfield known to be associated with elevated citation counts (40) – we tested whether this topic could confound the differences observed between flagged and non-flagged papers. Non-coding RNA papers, while representing 11% of the total dataset, accounted for 59% of flagged papers at the 0.6 threshold, confirming the disproportionate presence of this subfield among flagged papers. After restricting the analysis to papers mentioning non-coding RNA keywords (miRNA, lncRNA, circRNA) in their titles or abstracts, flagged papers still displayed higher citation and generally lower attention rate ratios than non-flagged ncRNA papers (SI Appendix, Fig. S4).

To supplement the rate ratio analysis, we investigated the association between citations and *Mendeley* readers at the article level (SI Appendix, Fig. S5). We note the strong positive association between increased readers and increased citations in the journal *Nature Cancer*, which had no flagged papers, highlighting the expected pattern of more readers leading to more citations. Across journals, flagged papers invariably received more citations than non-flagged papers for comparable numbers of *Mendeley* readers, while rarely reaching the highest reader numbers attained by non-flagged papers. These patterns held when considering only papers affiliated with Chinese institutions (SI Appendix, Fig. S6) and using online accesses in place of *Mendeley* readers (SI Appendix, Fig. S7). Together, these findings reveal a decoupling between citations and scientific attention, whereby flagged papers accumulate substantially more citations despite attracting fewer readers.

### Flagged papers cite and are cited by other flagged papers that cluster in specific journals

Having established that flagged papers received more citations than non-flagged papers but with fewer readers, we sought to understand the origins of this citation uptake and whether this advantage could reflect citation exchange. We applied our detection model (21) to the 2.4 million citing papers and the 1.5 million referenced papers using the 0.6 paper mill probability threshold. As our detection model was trained on cancer research papers, we performed a sensitivity analysis restricting both citing and referenced papers to the same set of 20 journals, to assess the impact of potential out-of-domain misclassification. We also performed a sensitivity analysis to control for potential citation homophily bias (41).

Flagged index papers were associated with higher proportions of both flagged citing and flagged referenced papers compared with non-flagged papers (Fig. 4, raw data points shown in SI Appendix, Fig. S8; SI Appendix, Tables S5-6), with a clear separation between the density distributions and centroids for most journals. An explanation for this pattern is that paper mills are producing many papers that cite their previous papers. Among flagged index papers, a higher baseline proportion of citing papers were themselves flagged, and this proportion increased in tandem with the share of flagged referenced papers. The centroid of non-flagged index papers was positioned in the lower range of the X-axis, with density contours remaining concentrated near zero – suggesting they receive citations from flagged papers without a corresponding increase in flagged referenced papers, as observed for *Nature Cancer*. The observed pattern holds at the 0.9 paper mill probability threshold (SI Appendix, Fig. S9) and when restricting the analysis to papers affiliated with Chinese institutions (SI Appendix, Fig. S10), which are overrepresented among papers citing and referenced by flagged index papers (SI Appendix, Tables S7-8).

**Fig. 4:**
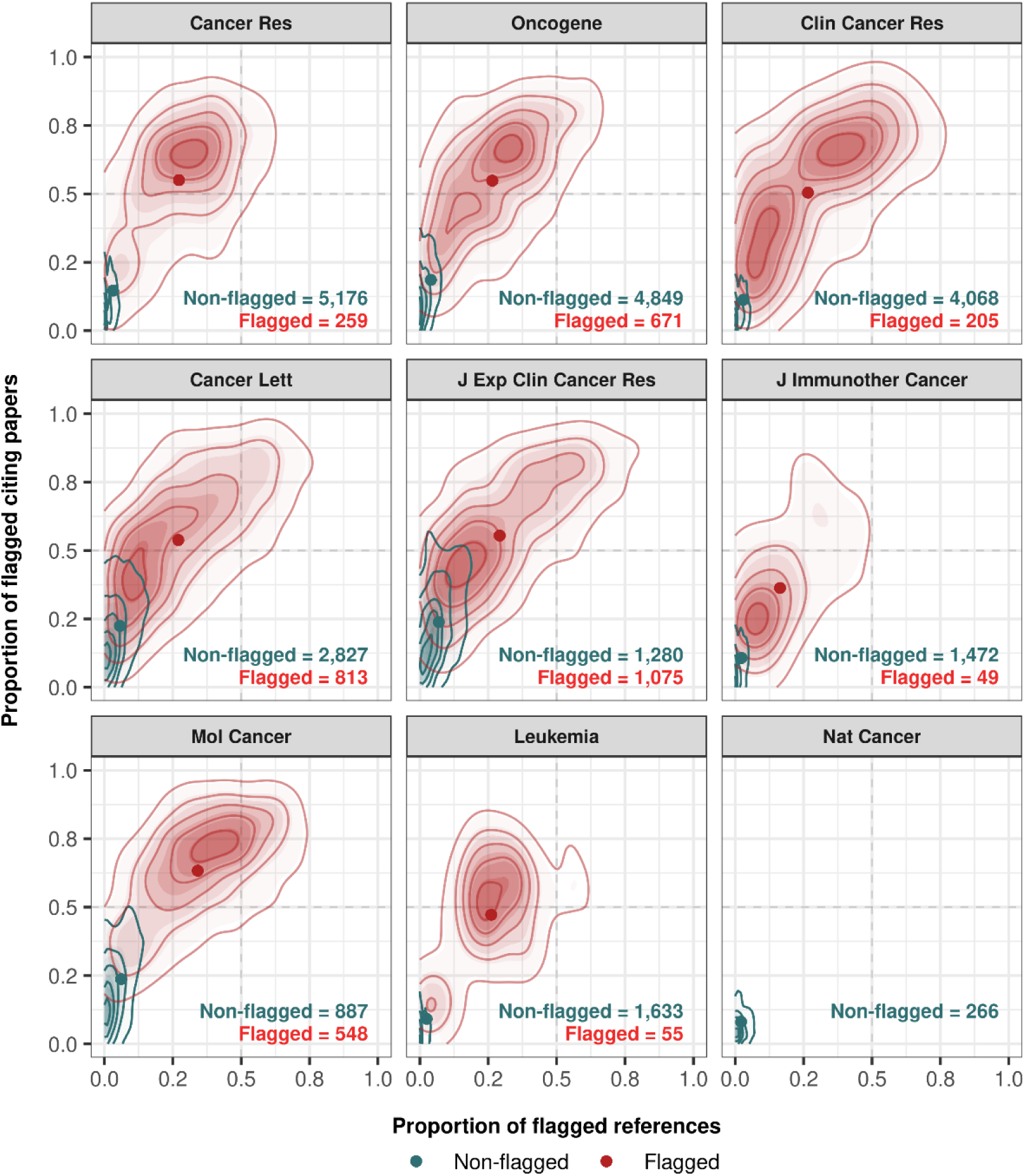
Two-dimensional kernel density estimates showing, for each of nine journals, the joint distribution of the proportion of references that cite flagged papers and the proportion of received citations that originate from flagged papers, separately for flagged and non-flagged index papers. All results used the 0.6 paper mill probability threshold. The first eight journals correspond to those with the largest numbers of papers in the dataset. Nature Cancer was included for comparison, as this journal had no flagged papers. Vertical and horizontal dashed lines indicate the midpoints of the X- and Y-axes (0.5). Contour lines show density levels from 10% to 90% of each group’s peak density, with points indicating each group’s centroid (mean). Journal names follow standard PubMed abbreviations. The distribution of individual data points underlying this figure is presented in SI Appendix, Fig. S8. The corresponding figure with index papers flagged at the 0.9 threshold is presented in SI Appendix, Fig. S9, and that restricted to China-affiliated papers is presented in SI Appendix, Fig. S10.

Using mixed-effects logistic regression, we found that citations to flagged index papers were strongly associated with much higher odds that the citing paper was also flagged (odds ratio = 6.5 with 95% CI: 6.3 to 6.8). References from flagged index papers were similarly associated with much higher odds that the referenced paper was also flagged (odds ratio = 10.1 with 95% CI: 9.7 to 10.6). These large odds ratios indicate strong clustering of flagged papers. Both associations remained robust after additional adjustment for Chinese authorship (odds ratio = 5.7 with 95% CI: 5.6 to 5.9 for citing papers and 7.9 with 95% CI: 7.6 to 8.2 for referenced papers). We finally assessed whether flagged papers draw citations from a more concentrated pool of sources using the Gini coefficient (SI Appendix). Flagged index papers showed higher inequality in both their citation and reference distributions (citation Gini 0.36 vs 0.19; reference Gini 0.35 vs 0.28), consistent with citations concentrating in a recurring set of papers.

In a first sensitivity analysis, restricting both citing and referenced papers to the same 20 journals further increased odds ratios to 9.5 (95% CI: 9.0 to 10.2) for citing papers and 14.3 (95% CI: 13.5 to 15.3) for referenced papers, indicating that potential out-of-domain misclassification for flagged papers does not drive these results. In a second sensitivity analysis, we introduced a proxy for research topic as an independent variable in the mixed-effects logistic regression model (see SI Appendix for details). The odds ratios remained elevated, indicating that the citation clustering effects we observe are unlikely to reflect topic homophily (citation model odds ratios: 4.5 to 5.7; reference model odds ratios: 7.1 to 9.3).

Early citations to flagged index papers predominantly originated from other flagged papers (SI Appendix, Fig. S11), with more than 75% of citations received during the first three years attributable to flagged papers at the 0.9 threshold (approximately 65% at the 0.6 paper mill probability threshold). This percentage declined progressively, approaching 50% after 10 to 12 years, while remaining stable at 10 to 20% throughout for non-flagged papers. The share of citations originating from review articles also increased markedly for flagged index papers from 2018 to 2024, reaching nearly 40% in 2024, compared with a stable 32 to 37% for non-flagged papers over the same period (SI Appendix, Fig. S12). These patterns suggest that the citation advantage of flagged papers is built early through contributions from other flagged papers, with literature reviews contributing more citations in recent years.

To further characterise citation flows at the journal level, we examined which journals were associated with citations to and references from flagged index papers using residuals from a Pearson chi-squared test (Fig. 5A, 5B). A subset of journals including *Biomedicine & Pharmacotherapy, Molecular Therapy - Nucleic Acids, Bioengineered* and *Pathology – Research and Practice* were in the upper-right quadrant – indicating that flagged index papers both cite and are cited by papers from these journals. Several of these journals have documented histories of integrity concerns, including delisting from bibliometric databases, paper mill targeting, and retraction campaigns (SI Appendix, Table S9). Three journals from the set of 20 molecular oncology journals (Fig. 2) – namely *Molecular Cancer*, *Journal of Experimental & Clinical Cancer Research*, and *Cancer Letters* – were also associated with citations to and references from flagged index papers.

**Fig. 5:**
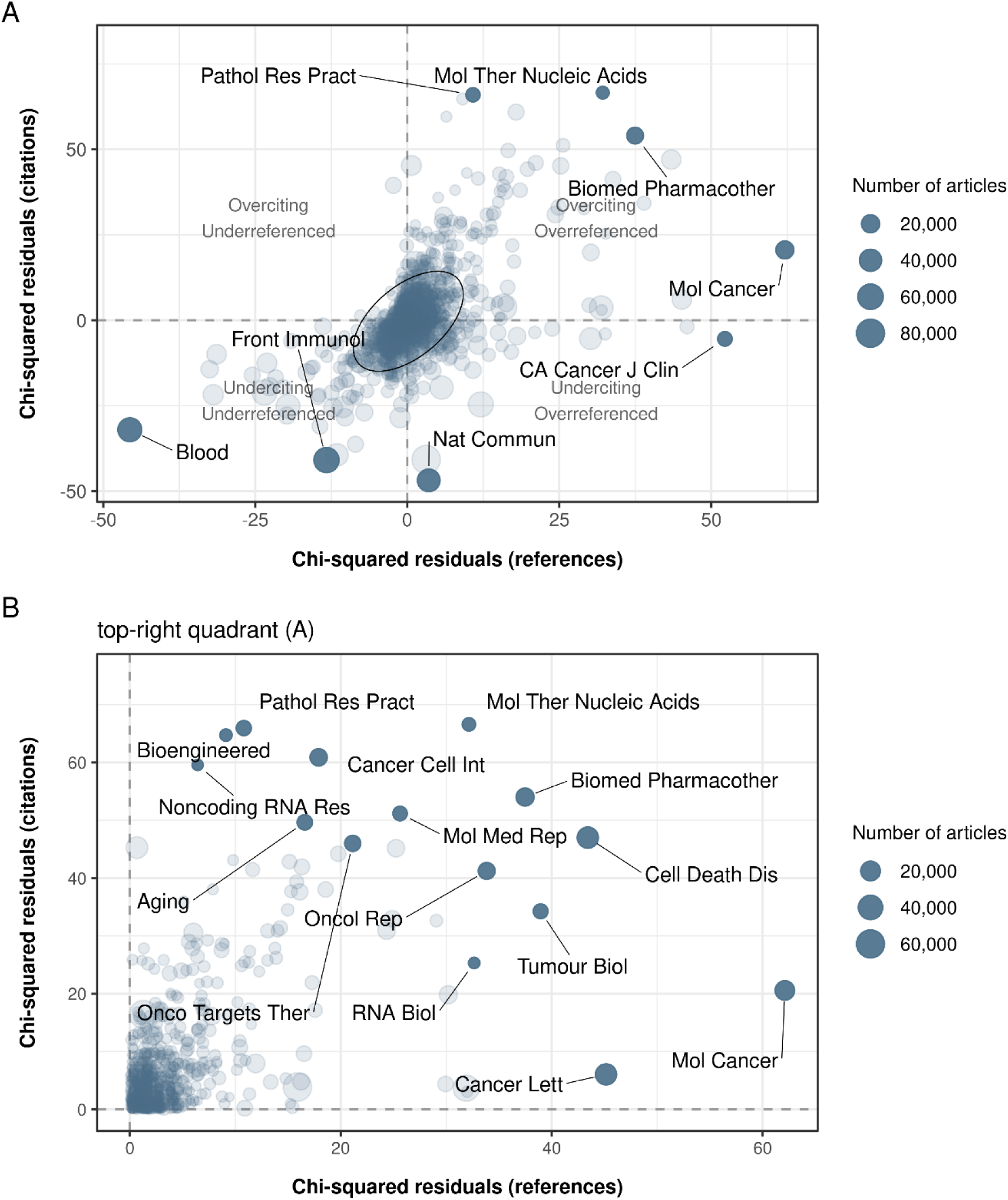
Association of journals with citations to (Y-axis) and references from (X-axis) flagged index papers, as determined by Pearson chi-squared residuals at the 0.6 paper mill probability threshold. Each point represents a journal, sized proportionally to the total volume of citations and references linked to flagged index papers. Journal self-citations and self-references were excluded. Journal names follow standard PubMed abbreviations. **(A)** All journals positively and / or negatively associated with citations to, and references from, flagged index papers are shown. The central ellipse shows a dense area covering 95% of journals where both residuals are relatively close to zero; grey points represent all remaining journals. **(B)** Journals showing excess citations to and references from flagged index papers (**(A)**, top-right quadrant). Labelled points correspond to the ten journals most overrepresented in citations and the ten most overrepresented in references from flagged index papers; some journals are in both groups. This pattern holds at the 0.9 paper mill probability threshold (SI Appendix, Fig. S13).

Some journals, such as *CA: A Cancer Journal for Clinicians*, were overrepresented in the reference lists of flagged index papers but underrepresented among papers citing them; a possible explanation for this pattern is that paper mills might cite papers from this high-profile journal to add legitimacy, but that papers in this journal do not more frequently cite paper mill publications. Conversely, other journals, including *Blood* and *Frontiers in Immunology*, showed a strong negative association with both citations to and references from flagged papers. Notably, very few journals were overrepresented in citations to flagged index papers without also being overrepresented in their reference lists. This reciprocal pattern supports the existence of a deliberate, non-random citation exchange, unlikely to arise under the null hypothesis of independence between journal identity and flagged status (both p-values were < 0.001 in the Pearson chi-squared test). Similar results were observed at both paper mill probability thresholds (SI Appendix, Fig. S13). Together, these findings show that flagged papers preferentially cite and are cited by other flagged papers, with these exchanges occurring rapidly after publication and forming self-reinforcing citation patterns that circulate through a subset of journals, consistent with an organised system.

### Flagged papers inflate citation metrics

To assess the extent to which flagged papers could influence journal-level citation metrics, we first quantified the number of citations received by flagged and non-flagged papers across the 20 journals. Flagged papers accounted for a substantial share of total citations in several journals (SI Appendix, Table S10), in some cases exceeding half of all citations received. At the 0.6 paper mill probability threshold, this percentage reached 57% for *Molecular Cancer* and *Journal of Experimental & Clinical Cancer Research*, 32% for *Cancer Letters*, and 17% for *Oncogene*. The same pattern held at the 0.9 probability threshold, although with lower overall percentages.

To further evaluate how citation flows to flagged papers could contribute to journal citation metrics, we calculated a metric analogous to the impact factor – the 2-year citation rate (2YCR) – for each year from 2014 to 2023, with and without flagged index papers (see SI Appendix for details). We did not calculate the familiar journal impact factor, as the definition of what constitutes a citable item in Web of Science is not fully transparent.

We focused on the six journals showing the largest average contribution of flagged index papers to the 2YCR – *Journal of Experimental & Clinical Cancer Research*, *Molecular Cancer*, *Oncogene*, *Cancer Letters*, *Neoplasia*, and *Cancer Research* – to illustrate the distortion that flagged papers introduce into journal citation metrics (Fig. 6). Full 2YCR contribution estimates with bootstrap 95% confidence intervals for all 20 journals across all years are provided in Dataset S2. The same visualisation for the 0.9 paper mill probability threshold is shown in SI Appendix, Fig. S14.

**Fig. 6:**
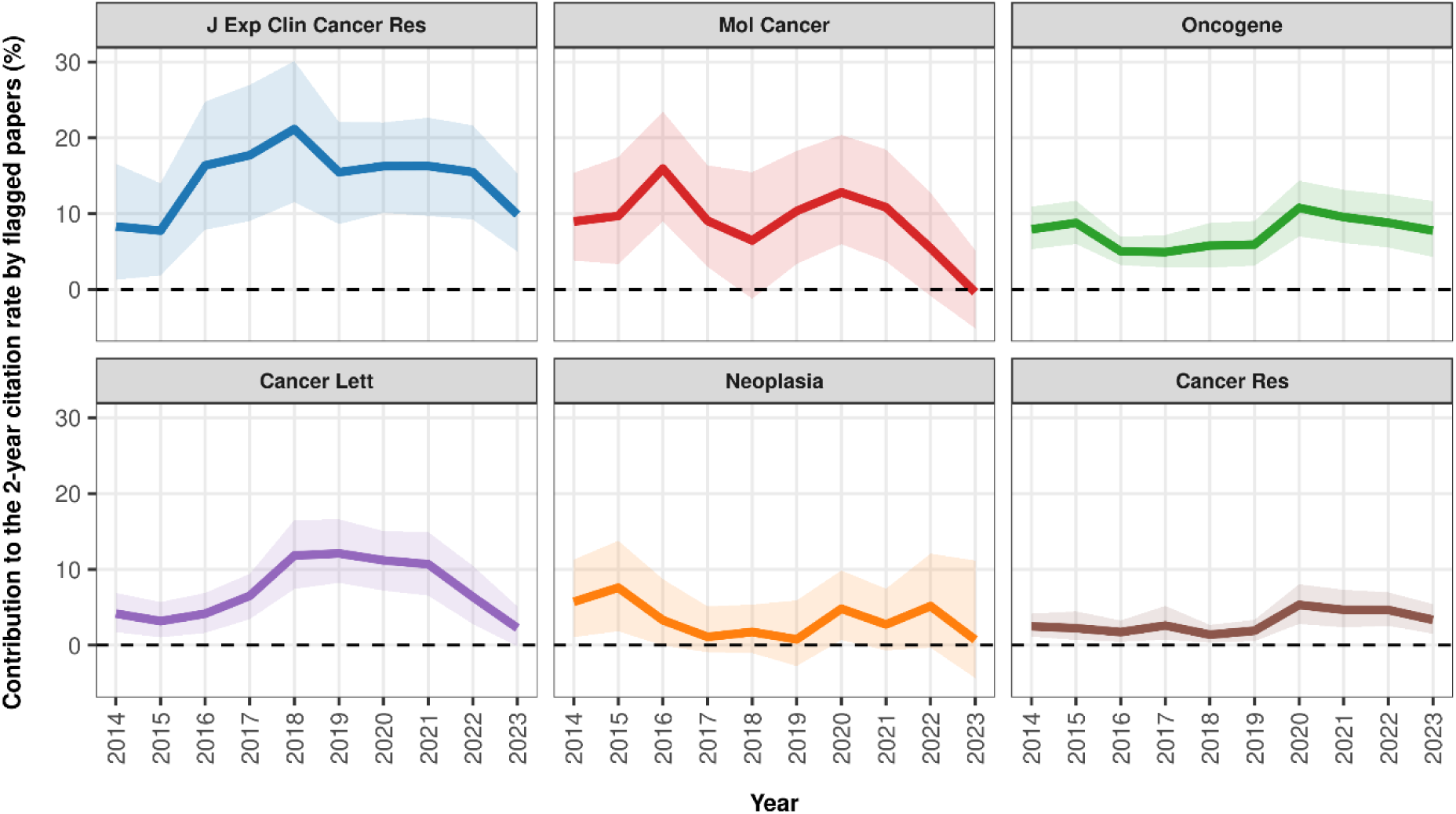
Contribution (%) of flagged index papers to the 2-year citation rate of 6 journals (Y-axis) at the 0.6 paper mill probability threshold, for each year of calculation (X-axis). For example, a value of 5% indicates that the journal’s 2-year citation rate would have been approximately 5% lower in the absence of flagged papers and their citations. Values below zero would indicate years in which flagged papers contributed less to the 2-year citation rate than non-flagged papers. 95% confidence intervals are shown as shaded bands around the lines. The equivalent visualisation for the 0.9 threshold is shown in SI Appendix, Fig. S14.

All six journals had positive contributions from flagged index papers to their 2YCR, although the gains varied considerably across journals and years. The relatively small number of papers published by each journal per year sometimes resulted in wide confidence intervals. Contributions from flagged index papers were highest for the *Journal of Experimental & Clinical Cancer Research* (10 to 17%, peaking at 21% in 2018) and *Molecular Cancer* (6 to 13%, peaking at 16% in 2016). A general decline in contributions is visible across most journals from 2021 to 2023, and confidence intervals are widest in 2023. While large confidence intervals reflect genuine uncertainty for journals or years with fewer contributing papers, contributions for the most affected journals remain positive across multiple years. These findings show that citations to flagged papers can inflate a journal’s citation metrics.

## Discussion

This forensic bibliometric analysis examines papers exhibiting textual similarity to paper mill papers in highly ranked molecular oncology journals. These flagged papers received 50 to 100% more citations than non-flagged papers, particularly in the early years after publication, yet attracted reduced reader attention. This pattern is counter-intuitive, as papers with more citations should have more readers. Citations typically accumulate progressively over time (42), and are strongly correlated with attention from the scientific community (42, 43).

No straightforward explanation accounts for this decoupling between attention and citations. A possible explanation is that paper mill-generated papers are not being widely read and are instead gaining citations through the organised behaviour of paper mills. Paper mill customers might pay for citations alongside their manuscript (44), or paper mills may systematically cite their products. Either way, any citation surplus may then trigger amplification from genuine researchers, as already highly cited articles tend to attract further citations through preferential attachment (42, 45). The increasing share of citations to flagged papers originating from review articles from 2018 to 2024 could further accelerate citation amplification, as literature reviews generally cite many papers.

Further evidence of organised citation exchanges comes from the citation flows surrounding flagged index papers, where a large fraction of citations and references originate from other flagged papers. The disproportionate representation of Chinese institutions among both flagged index papers and their citing papers is consistent with prior evidence linking paper mill activity to the Chinese academic environment (21, 46, 47). The cluster of journals simultaneously overrepresented in citations to and references from flagged index papers (Fig. 5) is consistent with an organised pattern of behaviour, with several of these journals having been reported as targets of paper mill activity or research fraud (SI Appendix, Table S9).

The fact that flagged index papers – which represent a minority of papers – can materially boost the journal citation metrics of the highest-impact molecular oncology journals raises concerns about the integrity of journal impact factors (48). Flagged papers were found across the first decile of molecular oncology journals (according to SCImago), ranging from *Molecular Cancer* (ranked #1) through *Experimental Hematology & Oncology* (#13) to *Oncogene* (#20) – indicating that paper mill activity is not confined to low-quality or predatory journals. These findings raise the possibility that paper mill activity has played a role in shaping journal impact factors – and by extension the perceived prestige – of some of the highest-ranked journals in molecular cancer research.

Flagged index papers were clustered by journal (Fig. 2), which is consistent with evidence that paper mills tend to focus on journals where they have published in the past (9). Most journals have incentives to publish highly cited papers, as this boosts their impact factor (49), which in turn increases submission numbers (50) and, in open-access models, earnings from article processing charges (51). This creates a conflict of interest for journals with quantity prioritised over quality and a reluctance to retract highly cited papers even when concerns are raised (52). There may even be a “win-win” scenario in which journals benefit from paper mill-engineered citations while paper mills supply citations to their clients. As highly cited papers disproportionately influence subsequent research (25), artificially inflated paper mill outputs may misdirect future research and, in molecular oncology, reduce the translation of preclinical findings into practice.

### Limitations

Several limitations relate to the paper mill detection model (21). It is a statistical model and will have misclassified individual papers; it provides no direct evidence of misconduct, and all conclusions rest on indirect supporting evidence. Citation motivations cannot be determined, precluding direct demonstration of paper mill intent. The model was trained on papers labelled as “paper mill” in the Retraction Watch database – a staff-assigned categorisation that will introduce some noise. It was, however, validated against an independent list of potential paper mill papers and the model performed comparably on both sources. Model predictions may be less stable on non-original-article publication types – a limitation that was expected to affect flagged and non-flagged groups equally.

The use of retracted papers as ground truth introduces circularity, as the identified corpus is disproportionately drawn from journals with historically high retraction rates. As retraction decisions are often delayed and can occur in batches, temporal coverage of training data is inherently uneven. Our findings may therefore not generalise beyond molecular oncology, and this approach may not capture paper mill products that evade retraction – whether through improved fabrication, by publishing in journals unlikely to retract, or by adapting methods that circumvent retraction-based detection.

While we adjusted for research topic within molecular oncology, sub-classifying molecular oncology papers is methodologically challenging – high-impact papers frequently examine multiple cancer hallmarks simultaneously, making clean topical boundaries difficult to define. Our embedding-based classification therefore does not exclude a topic homophily hypothesis. Furthermore, our analysis is restricted to the top decile of molecular oncology journals, and patterns may differ in lower-impact venues.

Some of the observed patterns may reflect structural and cultural features of the research environment in China rather than misconduct or paper mill activity. As a majority of citations to flagged index papers also come from Chinese authors, this could indicate “home-bias” citation patterns (53). This explanation is however not supported by the following findings: (i) adjusting for Chinese affiliation in the negative binomial models did not alter the observed patterns, either in citation or in attention; (ii) a majority of the papers citing flagged China-affiliated papers are themselves textually similar to paper mill papers; and (iii) these citing papers are clustered within problematic journals. We also acknowledge that *Mendeley* is less widely adopted in China, potentially underrepresenting Chinese researchers’ reading activity. However, Chinese authorship was explicitly adjusted for in both citation and attention models, and this limitation is expected to affect flagged and non-flagged papers comparably.

### Perspectives and future work

Citation network modelling could offer complementary insights, particularly for characterising citation clusters among flagged papers; future work could compute formal assortativity coefficients within the flagged subgraph and examine clustering patterns using citation graph topology. Similar analyses could be conducted in other domains including mathematics, computer science or materials science, where paper mill activity has been reported (44).

Our results indicate potential heuristics for detecting paper mill papers, notably the combination of high citations with relatively few readers – a pattern unlikely to arise for genuine papers – alongside a high early-citation inflation index (early citations / total citations) or journal-normalised citation surplus (paper citations / mean journal citations at matched time points), all of which could be formally tested in further research. These could be used to trigger more detailed research integrity investigations when coupled with other signs of potential misconduct. Reference lists may offer an additional first-line screening tool: unlike citations, which accumulate over time and may be diluted by citations from genuine papers, reference lists are finite, controlled by authors, and must efficiently serve the paper mill’s citation strategy – making them a more concentrated and hence detectable marker of manipulation. We believe that these metrics could represent promising screening indicators for publishers, indexing services, funders, and researchers.

Finally, the fact that systematic citation metric inflation may occur in high-impact journals should concern the scientific community and prompt concerted action. Citations are not just academic currency; they structure the flow of ideas, signal what is worth reading, and shape the directions that fields pursue. Their manipulation therefore corrupts not just individual metrics, but the collective process by which science builds on itself. Citations also carry direct financial weight – influencing article processing charges (APCs), funding decisions, and career trajectories – making their manipulation a profitable endeavour. Without oversight, this phenomenon may accelerate the selection of bad science – and ultimately of bad actors.

## Materials and Methods

### Definition of molecular cancer research

This research focused on molecular cancer research papers. We used the following definition, which guided journal and article selection: “*Molecular cancer research is focused on understanding how cancers develop, progress, and respond to treatment at the molecular and cellular levels. It studies the biological mechanisms that drive cancer by examining processes involving DNA, RNA, proteins, metabolites, and cellular signalling networks and how these processes regulate cell behaviour such as proliferation, survival, invasion, immune evasion, and metastasis*”. We focused on the highest-ranked cancer journals that publish molecular research, which are referred to as molecular oncology journals for readability. “Molecular cancer research” and “molecular oncology research” were considered equivalent.

### Journal selection and molecular cancer research filtering

Cancer journals in the top decile (D1) of the pooled 2024 SCImago (37) “Oncology” and “Cancer Research” rankings, ranked by citations per document (2 years), were retained and screened for relevance to molecular cancer research using their “Aims & Scope” section (see Dataset S1 for full journal selection process). All papers published in these journals between 2012 and 2023 – a window chosen to ensure complete citation history in OpenAlex – were extracted from the 2025 PubMed database. Molecular cancer research articles were identified using a validated BERT (54) classification model (accuracy = 0.96), excluding non-original-article publication types. Journals with fewer than 100 molecular research papers over the period were excluded, leaving 33,159 papers across 20 journals (Box 1). These papers were referred to as *index papers*, in contrast to *citing papers* which cite them, and *referenced papers* that they cite. See SI Appendix for further details.

#### Box 1: Highest-impact cancer journals publishing molecular research (ranked by 2024 SCImago citations per document, 2 years)

1. Molecular Cancer (30.3), 2. Journal of Hematology & Oncology (26.1), 3. Cancer Cell (24.7), 4. Nature Cancer (14.3), 5. Cancer Communications (12.3), 6. Journal of Experimental & Clinical Cancer Research (11.3), 7. Cancer Discovery (10.5), 8. Cancer Letters (9.2), 9. Journal for ImmunoTherapy of Cancer (9.1), 10. Neuro-Oncology (8.8), 11. Cancer Research (8.7), 12. Clinical Cancer Research (8.5), 13. Experimental Hematology & Oncology (8.4), 14. Journal of Thoracic Oncology (8.0), 15. Neoplasia (8.0), 16. NPJ Precision Oncology (7.8), 17. Leukemia (7.6), 18. NPJ Breast Cancer (7.0), 19. Cancer Immunology Research (6.9), 20. Oncogene (6.5).

### Citation counts and attention metrics

Citation counts, annual citation counts, and author country of affiliation (first listed country) were extracted for each index paper using the OpenAlex (39) API on 20th February 2026. For citing papers, DOI, country of authors’ institutional affiliations, journal, title, abstract text, and year of publication were extracted from the OpenAlex API on 1st March 2026; the same fields were extracted for referenced papers on 17th March 2026. To quantify attention, *Mendeley* readership, retrieved via Elsevier’s PlumX API on 25th March 2026, was used as a measure of personal engagement with a paper. Conversely, online accesses, collected from publisher websites on 20-24th February 2026, were used as a complementary measure of web traffic activity and were unavailable for a subset of journals. Detailed information is available in SI Appendix.

### Screening for suspected paper mill papers

Suspected paper mill papers were identified using our previously developed BERT model (21), trained on titles and abstracts of 2,202 retracted paper mill papers from the *Retraction Watch* Database (with accuracy = 0.91; sensitivity = 0.87; specificity = 0.96). Titles and abstracts of the 33,159 index papers were screened at two probability thresholds (0.6, corresponding to a 4% false positive rate on the model’s test set; 0.9, corresponding to a false positive rate below 1%). Citing and referenced papers were screened using the lower (0.6) threshold to assess whether flagged index papers also cite and are cited by suspected paper mill products. Papers predicted to share textual similarity with retracted paper mill papers were termed *flagged papers*, further distinguished as *flagged index*, *citing*, or *referenced papers*.

### Statistical models and sensitivity analyses

Negative binomial regression models (rate ratios, 95% CI) were adjusted for journal and year of publication, with cluster-robust standard errors (HC1, journal-by-year clusters). Models were not adjusted for the open-access status of papers, as this variable was largely absorbed by the journal fixed effect – the journals with the most flagged papers (e.g., *Molecular Cancer*, *Journal of Experimental & Clinical Cancer Research*) being fully open-access. Models were fitted on n = 33,159 index papers (4,085 / 29,074 flagged / non-flagged at the 0.6 threshold; 933 / 32,226 at 0.9), n = 33,149 for *Mendeley* readers, and n = 25,196 for online accesses. Sensitivity analyses additionally adjusted for Chinese authorship (n = 6,999 China-affiliated papers), restricted models to n = 3,524 non-coding RNA papers (miRNA, lncRNA, circRNA keywords), and reproduced models using Dimensions.ai citation counts instead of OpenAlex.

Mixed-effects logistic regression models (odds ratios, 95% CI) were adjusted for journal and year of publication, with a random intercept per index paper, and fitted on 2,389,626 citing and 1,538,819 referenced papers (journal self-citations / references retained, < 1% of records). Sensitivity analyses additionally adjusted for Chinese authorship, restricted citing and referenced papers to the same 20 journals as index papers (192,214 citing; 231,248 referenced papers) and introduced a research topic covariate derived from clustering of BERT embeddings (SI Appendix).

Journal-level overrepresentation of citations to and references from flagged index papers was assessed using Pearson chi-squared tests on journal-by-flagged-status contingency tables (self-citations / references excluded), with standardised residuals used to rank journals.

Trend lines were fitted using simple linear regression. For journal-panel visualisations, the eight journals with the largest numbers of papers were selected, with *Nature Cancer* as a negative control. The 5% most extreme values were excluded from Figs. S5-7 in SI Appendix (full data for SI Appendix, Fig. S5 in Fig. S15). A summary of all statistical models is in SI Appendix, Table S11. Figures were plotted using the R ggplot2 library (55) and most tables were plotted using the R gt library (56).

## Author contributions

JAB, BS and AGB conceptualised the study. BS, AGB, DC and JAB developed the methodology. BS and AGB developed and executed the code. BS, AGB, DC and JAB performed the analysis. BS and AGB wrote the manuscript. BS, AGB, JAB and DC reviewed and edited the manuscript. JAB and AGB secured the funding. AGB, JAB and DC supervised the work. All work reported in the paper was performed solely by the authors. BS is the guarantor of the study.

### Acknowledgements

The authors thank Matt Spick for his valuable review of this manuscript. They also acknowledge the members of the high-performance computing environment Tesla (*Institut de recherche mathématique de Rennes*, IRMAR, France) and Aqua (Queensland University of Technology, QUT, Australia) for granting access to their resources and for their technical support. The authors acknowledge the teams at PubMed, SCImago and PlumX for sharing their data. AI-assisted coding tools (Claude Code, Anthropic) were occasionally used to assist with code development. This paper was written using data obtained on 15th July 2026, from Digital Science’s Dimensions platform, available at https://app.dimensions.ai. Access was granted to subscription-only data sources under licence agreement.

## Data and code availability

Data used in this study are publicly available from PubMed annual XML datasets (https://ftp.ncbi.nlm.nih.gov/pubmed/baseline/), OpenAlex API and SCImago Journal and Country Rank. *Mendeley* readership data can be obtained from Elsevier’s PlumX. The fine-tuned models and code developed in our previous paper (21) are not publicly disclosed to prevent potential misuse by individuals seeking to evade fraud detection. Code related to the experiments in this paper can be accessed through this GitHub repository: BaptScc/mills-distort-citation-metrics.

