## Supplementary files for "Suspected distortion of citations in high-impact cancer journals"

### Supporting Information Text

#### Cross-checks against external integrity flags

PubPeer comments were collected by web scraping between 13th and 14th July 2026 for the 4,085 index papers flagged at the 0.6 threshold, and for a comparison subset of 4,088 non-flagged index papers sampled proportionally to each journal's number of non-flagged papers. Each retrieved comment was then classified as either raising a direct integrity concern or not, using GPT-4o (OpenAI API) through prompting. The full Problematic Paper Screener (PPS) report (1) was downloaded on 15th July 2026 and matched to index papers by DOI. For both sources, the proportion of index papers found to raise integrity concerns was computed and compared between flagged and non-flagged index papers.

#### Concentration of the citing and referenced paper pools

For each group of index papers (flagged and non-flagged) at the 0.6 threshold, we built a count vector of how often each distinct citing paper appeared across the group and computed the Gini coefficient (2) on that vector. The same was done for referenced papers. The Gini coefficient can be defined as follows:

$$G = \frac{2\sum_{i=1}^{n} i\, y_i}{n\sum_{i=1}^{n} y_i} - \frac{n+1}{n}$$

Where $y_i$ is the number of times the $i$-th distinct citing (or referenced) paper appears across the flagged or non-flagged group, sorted in ascending order, and $n$ is the total number of distinct citing (or referenced) papers in the group.

As the Gini coefficient depends on the number of index papers contributing to the pool, and non-flagged index papers outnumber flagged ones, the non-flagged coefficients were estimated from 20 random subsets of non-flagged index papers drawn without replacement and matched in size to the flagged group (4,085 index papers) and averaged across draws. Coefficients for the flagged group were computed directly on the full set.

**Controlling for topical homophily using BERT-based topic clusters**

One of the potential confounders that could explain the association between flagged index papers and flagged citing/referenced papers is topical homophily (3). As papers from a certain research field tend to preferentially cite other papers from their field, an over-representation of papers from a certain research domain (e.g., non-coding RNAs) could have influenced the odds ratios.

To attempt to control for this bias, in a sensitivity analysis, we produced embeddings from index papers' titles and abstracts using BERT (4) base (from the *google-bert/bert-base-uncased* Hugging Face repository). We extracted the 768-dimension vector from the last hidden state of BERT (CLS token) for each of the 33,159 index papers. To reduce dimensionality, we used the Principal Component Analysis method (PCA) with 50 components. The PCA components were then sub-classified into textual similarity clusters, using the K-means method (unsupervised clustering method, based on centroids). We believe that textual similarity is a good proxy for topic similarity, although we acknowledge that this approach has limitations. Both PCA and K-means methods were called from the *Scikit-learn* Python library (5). The number of K clusters was defined arbitrarily in a range of [5, 10, 20, 30, 50, 100] to assess the drift of odds ratios under finer classification.

Each K-cluster classification was used to potentially explain the link between citing / referenced papers and index papers in the mixed-effects logistic regression model, defined as:

$$RPS \ / \ CPS \sim IPS + journal + year + KCC + (1 \mid IPD)$$

Where RPS = Referenced Paper Status; CPS = Citing Paper Status; IPS = Index Paper Status; journal = journal of index paper; year = year of publication of index paper; KCC = K-Cluster Classification; IPD = Index Paper DOI. The parameter (1 | IPD) is a random intercept accounting for correlation between citing / referenced papers sharing the same index paper. The mixed-effects logistic regression model used the *lme4* R library (6).

The choice of BERT over manual keyword selection was motivated by the overlapping nature of topics within molecular oncology, and we felt a more comprehensive approach was better suited to this task. Although optimisation methods exist for the best number of K clusters, we report the citation / reference odds ratios across a range of cluster numbers to demonstrate that our results are not an artefact of any particular classification. Main themes for each K=10 cluster were derived from the most common unigrams and bigrams, as presented in the table below.

| Cluster | No. of papers | Main theme(s) |
|---|---|---|
| 1 | 2,627 | Resistance; combination therapy |
| 2 | 3,028 | Immune checkpoints; micro-RNAs |
| 3 | 4,215 | Signalling pathway; metastasis |
| 4 | 3,568 | Tumour biology |
| 5 | 5,220 | Micro-RNAs |
| 6 | 2,154 | Breast cancer |
| 7 | 3,946 | Treatment response; checkpoint inhibitors |
| 8 | 1,887 | Biomarkers; mutation |
| 9 | 2,995 | Immune micro-environment |
| 10 | 3,519 | Micro-RNAs; breast cancer |

**Journal citation metric calculation**

To study the contributions of flagged index papers to journal citation metrics, we calculated a journal citation metric, analogous to SCImago's citations per document (2 years), referred to as the *two-year citation rate*, defined as the number of citations received in year *i* by articles published in years *i - 1* and *i - 2*, divided by the number of published articles in the same two years. As *OpenAlex* provides annual citation data from 2012 onwards, our metric was computed for the years 2014 to 2023, 2014 being the earliest year for which the required two-year citation window could be fully reconstructed. The two-year citation rate was calculated as:

$$2YCR_i = \frac{\sum_{\{\alpha \in P_{i-1,\ i-2}\}} \mathrm{C}_{\alpha,i}}{\mathrm{P}_{\{i-1,\ i-2\}}}$$

Where $2YCR_i$ is the two-year citation rate of a journal for the year $i$, $C_{\alpha,\, i}$ is the number of citations received by paper $\alpha$ during the year $i$ and $P_{i-1,\, i-2}$ is the set of papers published in the years *i - 1* and *i - 2*.

The modified two-year citation rate, without flagged papers, was calculated as follows for both 0.6 and 0.9 thresholds:

$$2YCR_i^{(-F)} = \frac{\sum_{\{\alpha \in P_{i-1,\ i-2} \setminus F\}} \mathrm{C}_{\alpha,i}}{\mathrm{P}_{\{i-1,\ i-2 \setminus F\}}}$$

Where $F$ is the set of flagged papers (either at the 0.6 or 0.9 probability threshold), which is excluded from the calculation of the two-year citation rate.

For each journal, the contribution of flagged papers to the two-year citation rate was calculated as:

$$\text{Contribution (\%)} = \frac{2YCR_i - 2YCR_i^{(-F)}}{2YCR_i} \times 100$$

Bootstrap 95% confidence intervals (1,000 iterations) were computed for each journal and year separately, by resampling papers with replacement. Two-year citation rate values were compared with values from SCImago's citations per document (2 years) to assess agreement, showing a very strong correlation (Pearson correlation = 0.98) and a mean absolute error of 2.3 points. The absolute difference can be explained by differences in citation sources and in the types of documents included in the calculation of journal metrics.

**Journal selection and molecular cancer research filtering**

Journals appearing in both the SCImago "Oncology" and "Cancer Research" rankings were deduplicated (472 journals), of which the top 50 (D1) were retained. Review-specific journals (e.g., *Nature Reviews Cancer*) and non-cancer-specific journals (e.g., *Signal Transduction and Targeted Therapy*) were excluded. Aims & Scope screening was validated through a keyword analysis of the same section. A journal was considered to potentially publish molecular oncology research when at least one related keyword was found (e.g., "wet lab experiment"). This first step excluded 25 of the 50 initially selected journals.

All 149,215 papers published in the 25 remaining D1 cancer journals were extracted from the 2025 PubMed database using the journals' ISSNs, excluding empty abstracts and duplicated PMIDs or DOIs. The BERT (4) model used to identify molecular cancer research (accuracy = 0.96, sensitivity = 0.93, specificity = 0.97) was trained on 5,000 papers labelled using GPT-4o from the OpenAI API, and classified papers from their PubMed metadata, excluding clinical trials, reviews, comments, editorials, and other non-original-article types (correction and retraction notices, letters, etc.). Agreement between model predictions and a human expert (JAB) was evaluated on a sample of 100 papers (observed agreement = 0.89, Gwet's AC1 = 0.82, 95% CI 0.71 to 0.93). We restricted the analysis to journals publishing more than 100 molecular cancer research papers, resulting in a final set of 20 journals. Retracted original research articles were retained.

**Citation counts and attention metrics**

Annual citation counts from OpenAlex were used to calculate the number of early citations received by each index paper, 1 and 3 years after publication. The quality of the OpenAlex citation data was validated against Dimensions citation data (extracted on 15th July 2026) and achieved a Pearson correlation of 0.99 and a mean absolute error of 6.2 citations due to indexing differences in the two databases.

For each citing paper, of the 2,742,150 available citations, 2,596,936 (94.7%) were retrieved. Missing abstract texts were retrieved from the PubMed baseline using the papers' DOIs. Ultimately, 207,310 abstracts (8%) and 52,331 country affiliations (2%) remained unavailable. Similarly, of the 1,643,255 referenced papers, 1,548,987 were retrieved (94.3%) and 10,168 abstracts (<1%) and 22,966 country affiliations (1.5%) remained unavailable. All missing records were also excluded from the corresponding visualisations.

We initially considered Altmetric as a comprehensive proxy for attention, but did not use it as major Chinese social media platforms are not covered by Altmetric (7). We chose *Mendeley* (https://www.mendeley.com/) instead, an online reference manager that allows researchers to organise papers of interest, with 2.5 million users in 2013 (8) – that we believe to be used internationally. A paper's *Mendeley* readership count reflects the number of individual users who have added it to their personal library and therefore captures personal engagement with a paper.

We also used online accesses data, which typically reflect the number of times a paper's full text or abstract page was viewed or downloaded. We believe this metric is universal and captures worldwide readership activity distinct from *Mendeley* readers. It was often unclear whether the metric displayed on a journal page reflected page views, downloads, or a combination of both; however, this disparity was accounted for by adjusting for journal in the models. As Elsevier journals do not display access metrics, access counts could not be retrieved for *Cancer Cell*, *Cancer Letters*, *Journal of Thoracic Oncology* and *Neoplasia*. Access metrics were not available for the *Journal for ImmunoTherapy of Cancer*, and those for *Cancer Communications* were only available from 2022 onward. Given the small number of remaining *Cancer Communications* papers, the latter were excluded from the online access analysis.

#### Supplementary Figures

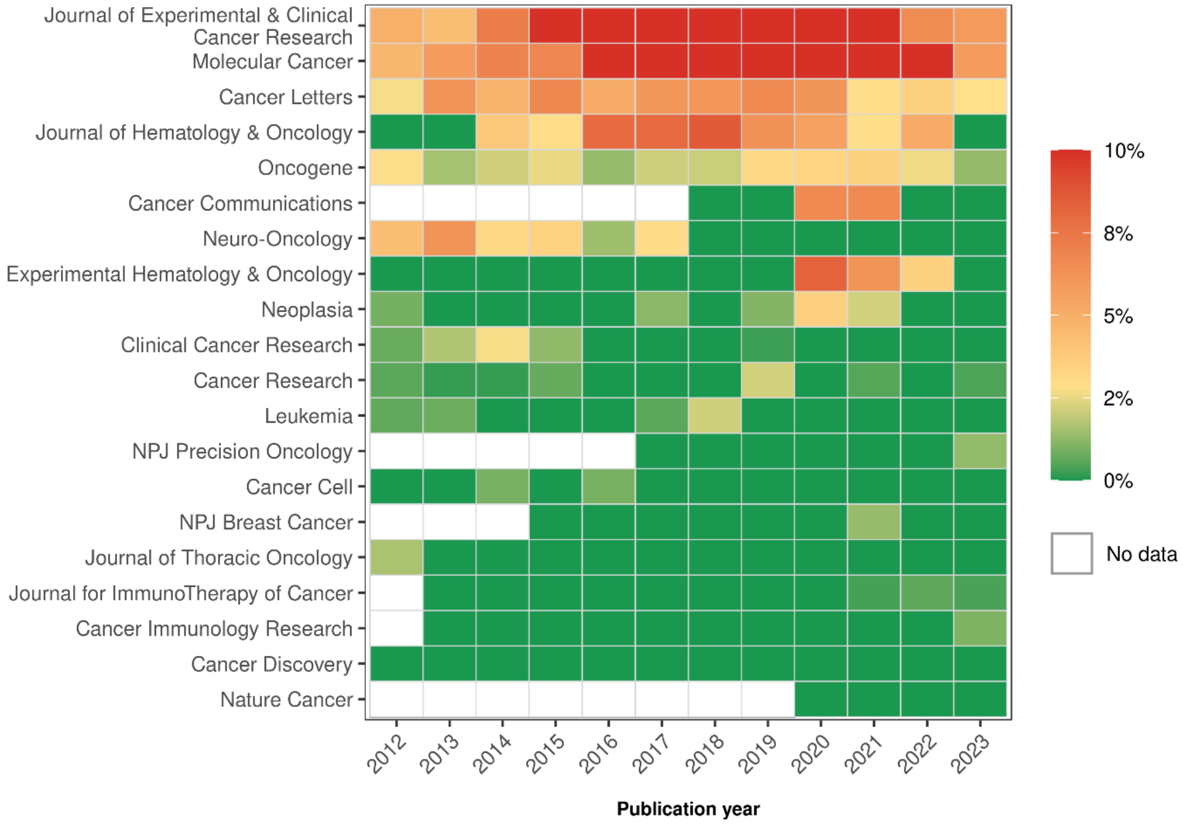


*Fig. S1: Heatmap showing yearly percentages of flagged papers across leading molecular cancer research journals (0.9 paper mill probability threshold). Journals are ordered according to their overall percentage of flagged papers during the study period. The colour scale is centred at 2.8%, which is the overall mean percentage of flagged papers. Blank cells indicate missing data, either because the journal had not been established or because no molecular cancer research papers were published in that year in that journal. Values are capped at 10%, with all percentages over 10% displayed at the maximum colour intensity.*

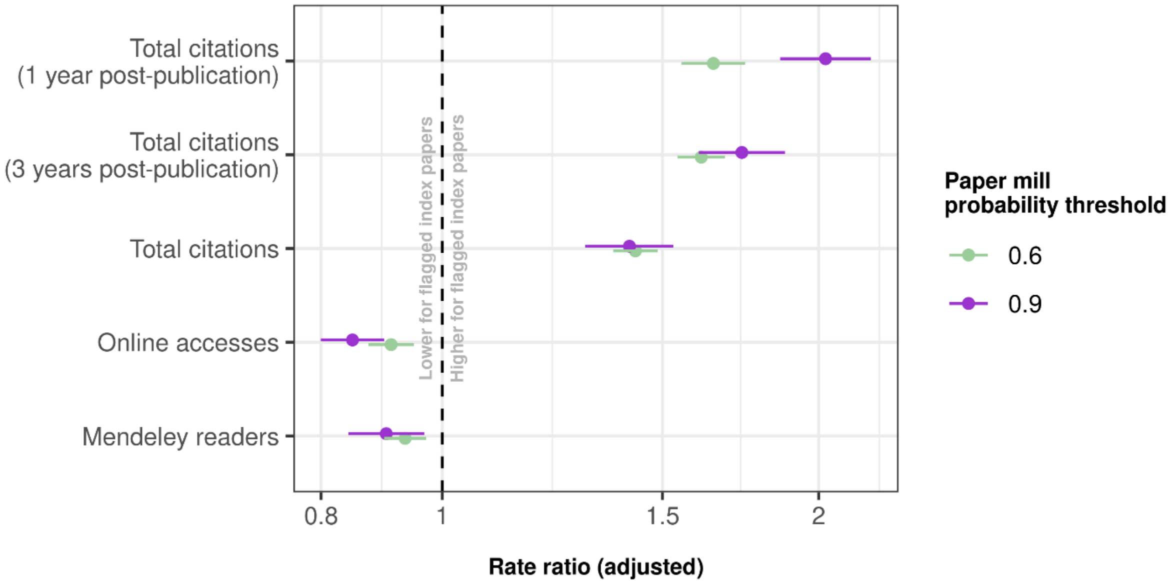


*Fig. S2: Adjusted rate ratios (RR) for citation and attention outcomes associated with flagged index papers. Attention outcomes (Mendeley readers and online accesses) reflect total cumulative attention since publication. Points represent adjusted rate ratios estimated from negative binomial regression models, comparing flagged and non-flagged index papers based on the paper mill probability threshold (0.6 and 0.9). Horizontal bars are 95% confidence intervals with cluster-robust standard errors (Journal×Year). Models were adjusted for journal, year of publication and the geographic origin of papers (Chinese versus non-Chinese authorship). Rate ratios are shown on a logarithmic scale. The vertical dashed line indicates a rate ratio of 1 (no difference between flagged and non-flagged papers).*

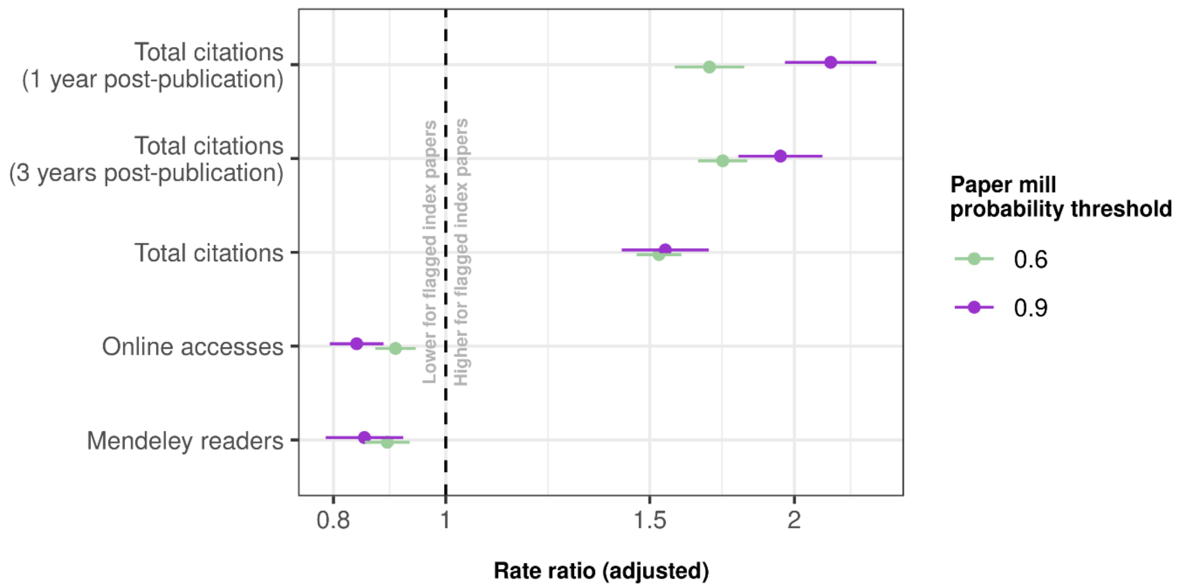


*Fig. S3: Adjusted rate ratios (RR) for citation and attention outcomes associated with flagged index papers, using citation data extracted from Dimensions.ai instead of OpenAlex as a sensitivity analysis to control for potential database-specific artefacts. Attention outcomes (Mendeley readers and online accesses) reflect total cumulative attention since publication. Points represent adjusted rate ratios estimated from negative binomial regression models, comparing flagged and non-flagged index papers based on the paper mill probability threshold (0.6 and 0.9). Models were adjusted for journal and year of publication. Horizontal bars are 95% confidence intervals with cluster-robust standard errors (Journal×Year). Rate ratios are shown on a logarithmic scale. The vertical dashed line indicates a rate ratio of 1 (no difference between flagged and non-flagged papers).*

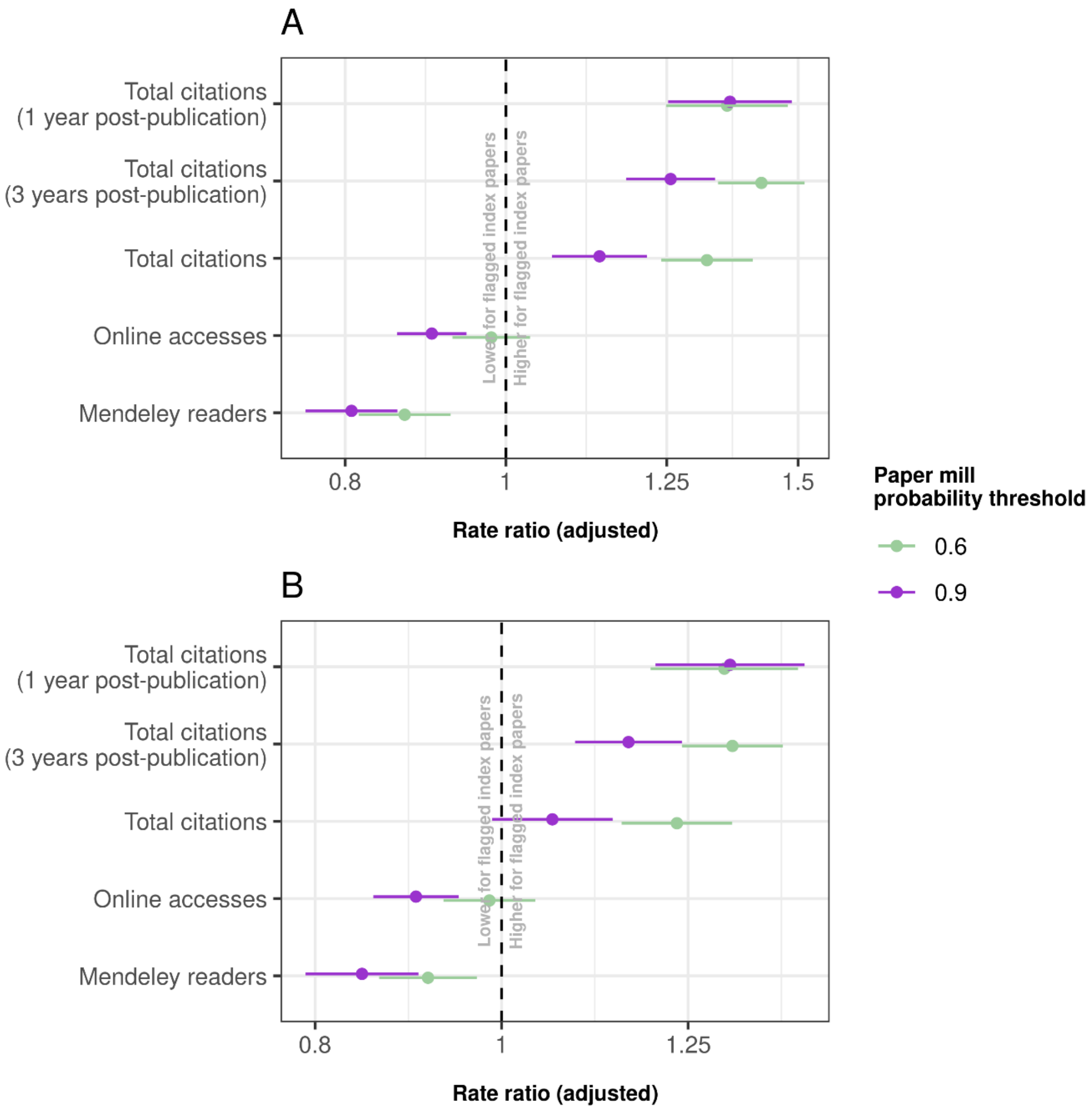


*Fig. S4: Adjusted rate ratios (RR) for citation and attention outcomes associated with flagged index papers. This analysis was conducted on the subset of papers associated with non-coding RNA research. Attention outcomes (Mendeley readers and online accesses) reflect total cumulative attention since publication. Points represent adjusted rate ratios estimated from negative binomial regression models, comparing flagged and non-flagged index papers based on the paper mill probability threshold (0.6 and 0.9). Horizontal bars are 95% confidence intervals with cluster-robust standard errors (Journal×Year). (**A**) Models were adjusted for journal and year of publication. (**B**) Models were further adjusted for the geographic origin of papers (Chinese versus non-Chinese authorship). Rate ratios are shown on a logarithmic scale. The vertical dashed line indicates a rate ratio of 1 (no difference between flagged and non-flagged papers).*

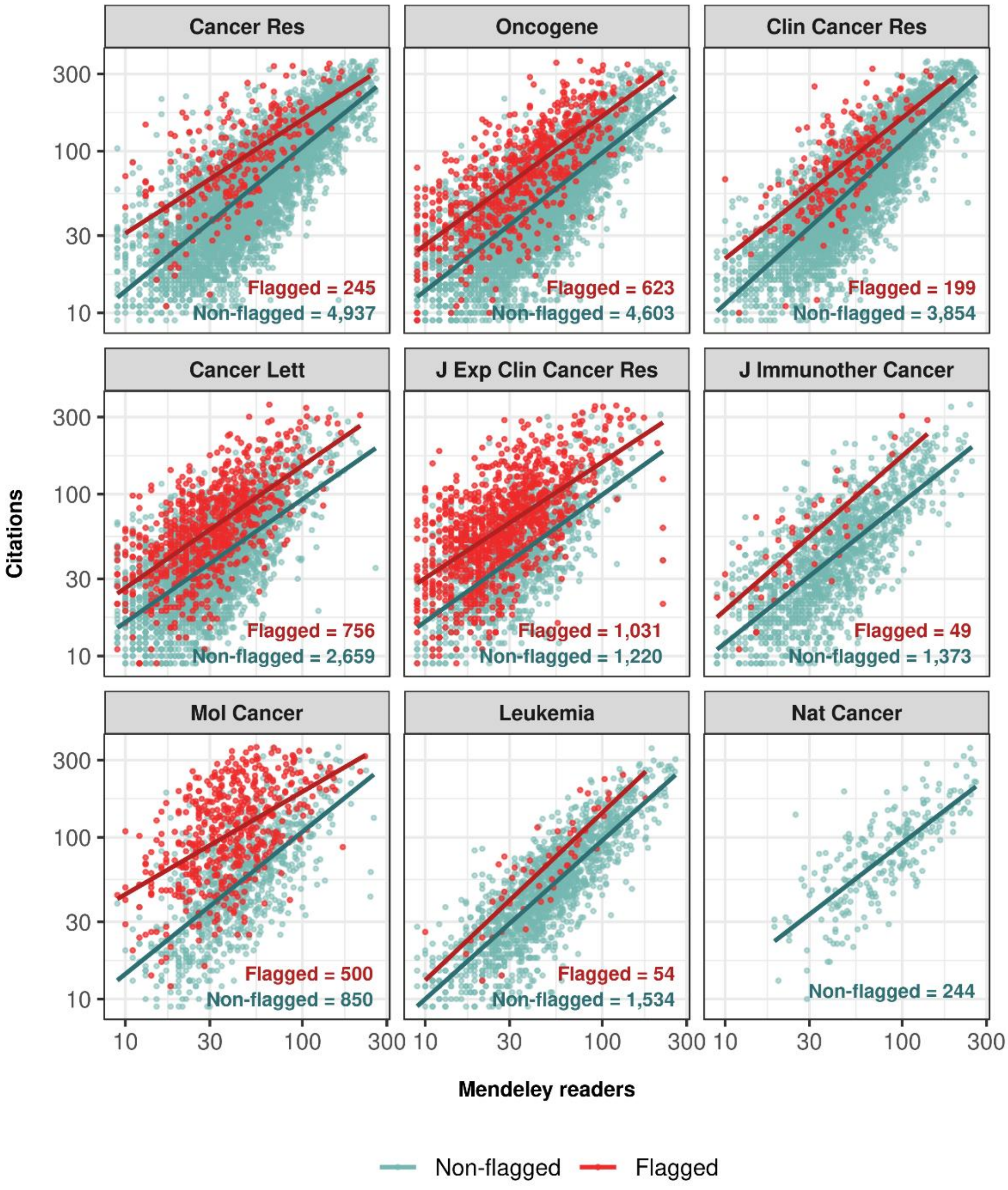


*Fig. S5: Association between citation counts and Mendeley readers across flagged and non-flagged papers in 9 journals at the 0.6 paper mill probability threshold. Both variables are presented on a log10 scale. The first eight journals correspond to those with the largest numbers of papers in the dataset. Nature Cancer was included for comparison, as this journal had no flagged papers. Each point represents an individual paper, and solid lines indicate fitted linear regression models stratified by flagged and non-flagged papers. For both variables, we plot the central 95% of values for visualisation purposes.*

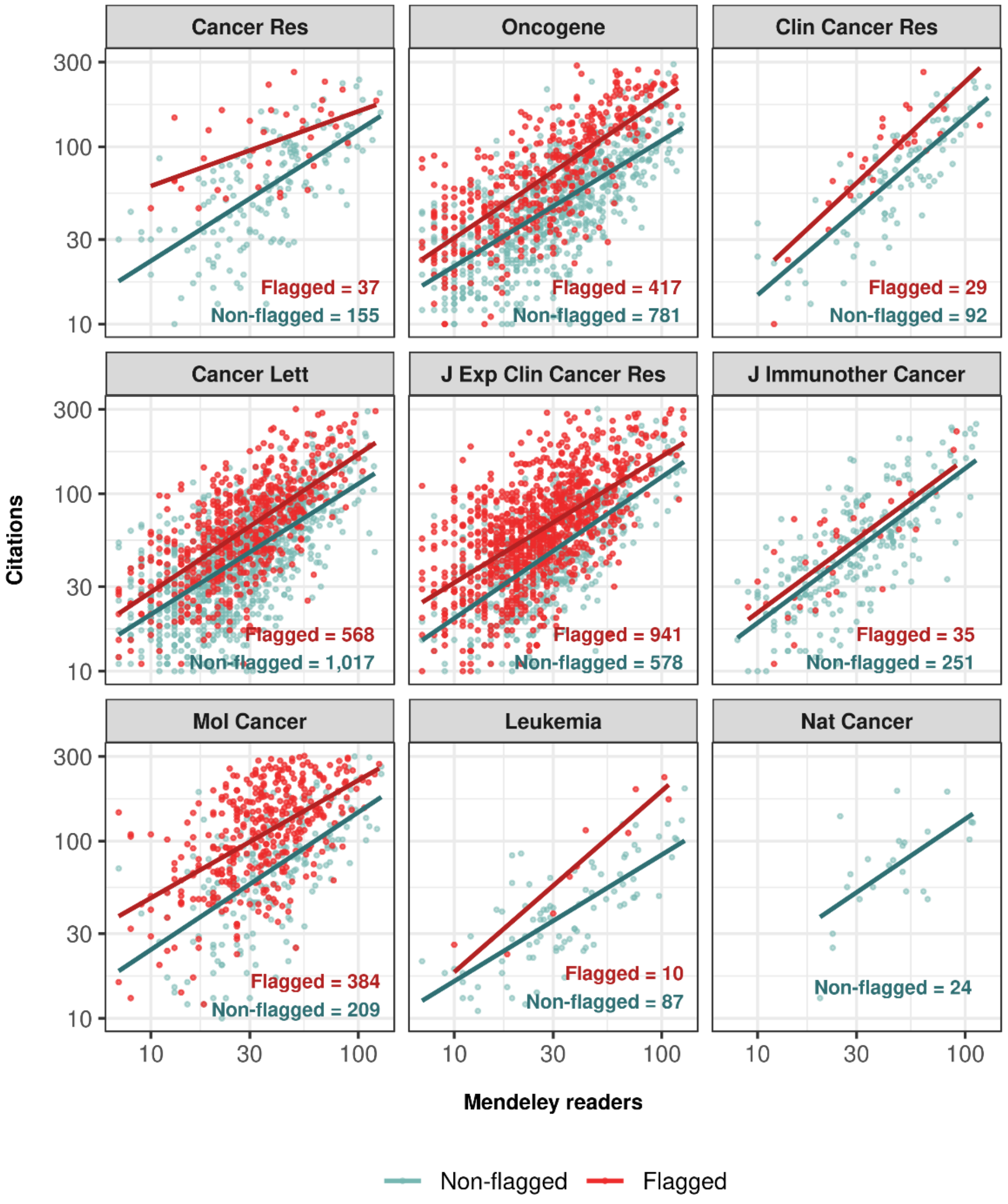


*Fig. S6: Association between citation counts and Mendeley readers across flagged and non-flagged papers in 9 journals at the 0.6 paper mill probability threshold. Only China-affiliated papers are shown. Both variables are presented on a log10 scale. The first eight journals correspond to those with the largest numbers of papers in the dataset. Nature Cancer was included for comparison, as this journal had no flagged papers. Each point represents an individual paper, and solid lines indicate fitted linear regression models stratified by flagged and non-flagged papers. For both variables, we plot the central 95% of values for visualisation purposes.*

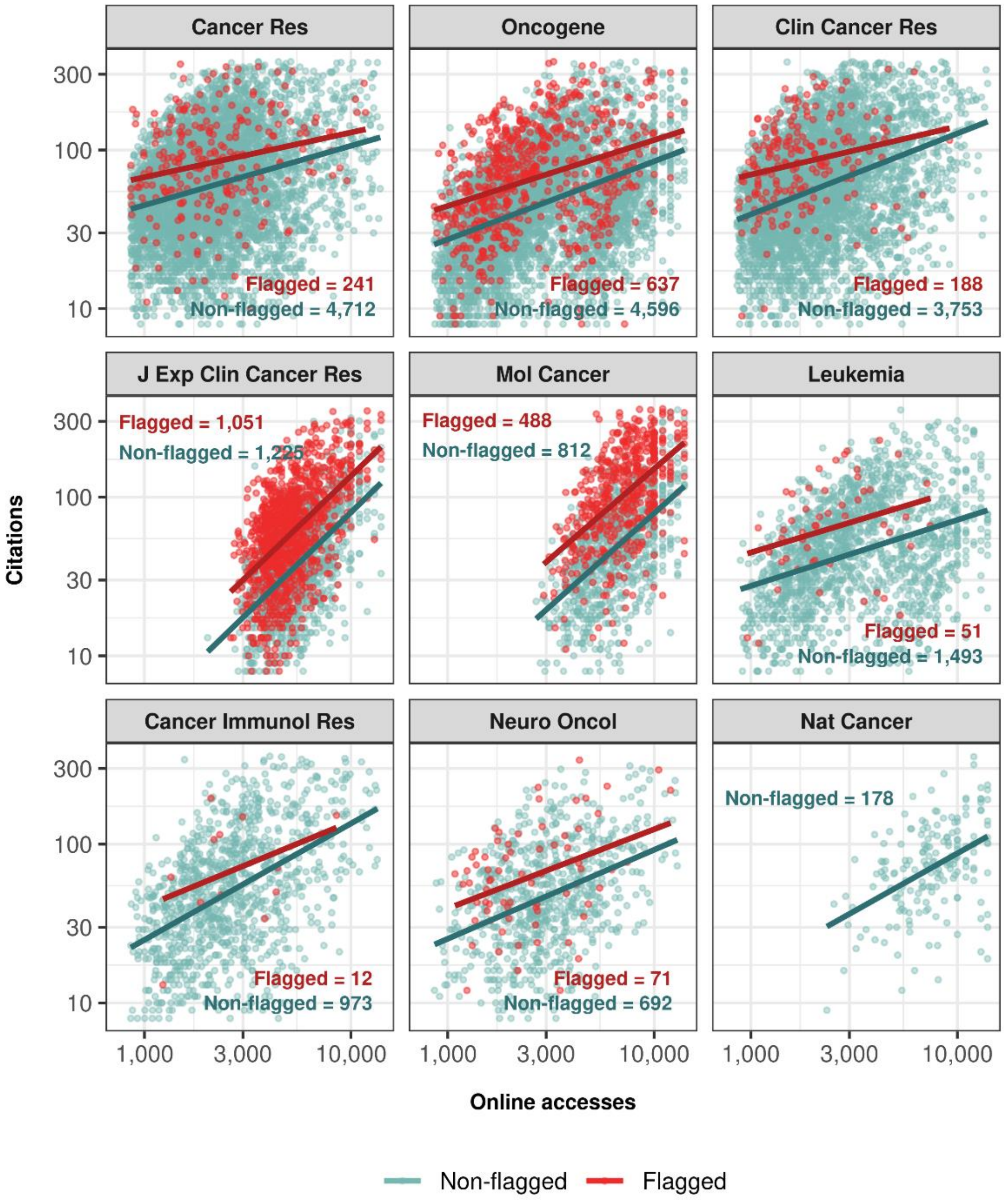


*Fig. S7: Association between citation counts and online accesses across flagged and non-flagged papers in 9 journals at the 0.6 threshold. Both variables are presented on a log10 scale. The first eight journals correspond to those with the largest numbers of papers in the dataset. Nature Cancer was included for comparison, as this journal had no flagged papers. Each point represents an individual paper, and solid lines indicate fitted linear regression models stratified by flagged and non-flagged papers. Values are rounded above 10 thousand accesses for Springer Nature journals. For both variables, we plot the central 95% of values for visualisation purposes.*

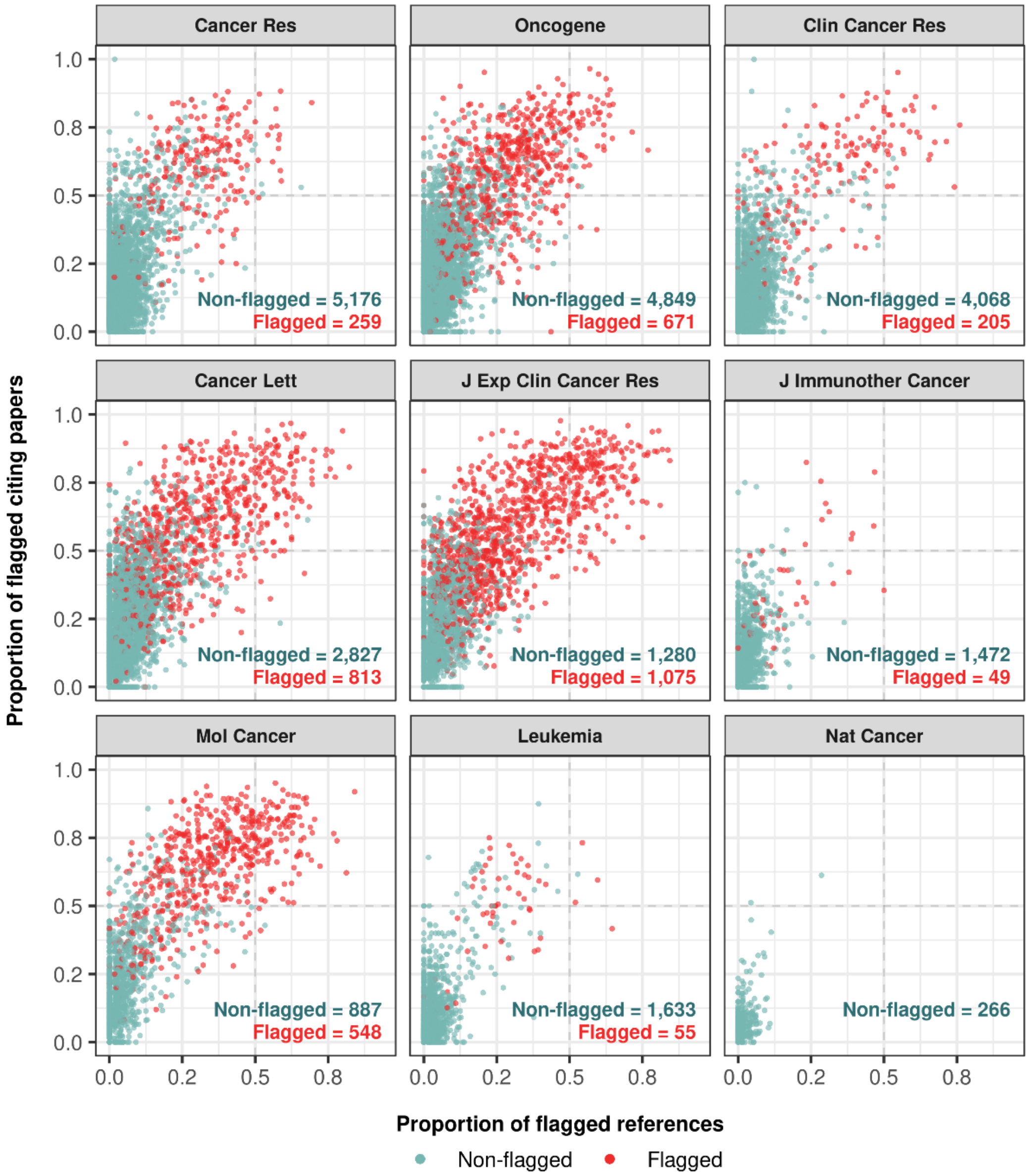


*Fig. S8: Scatterplot showing, for each index paper, the proportion of references that cite flagged papers and the proportion of received citations that originate from flagged papers across 9 journals. All results used the 0.6 paper mill probability threshold. The first eight journals correspond to those with the largest numbers of papers in the dataset. Nature Cancer was included for comparison, as this journal had no flagged papers. Vertical and horizontal dashed lines indicate the midpoints of the X- and Y-axes (0.5). Journal names follow standard PubMed abbreviations.*

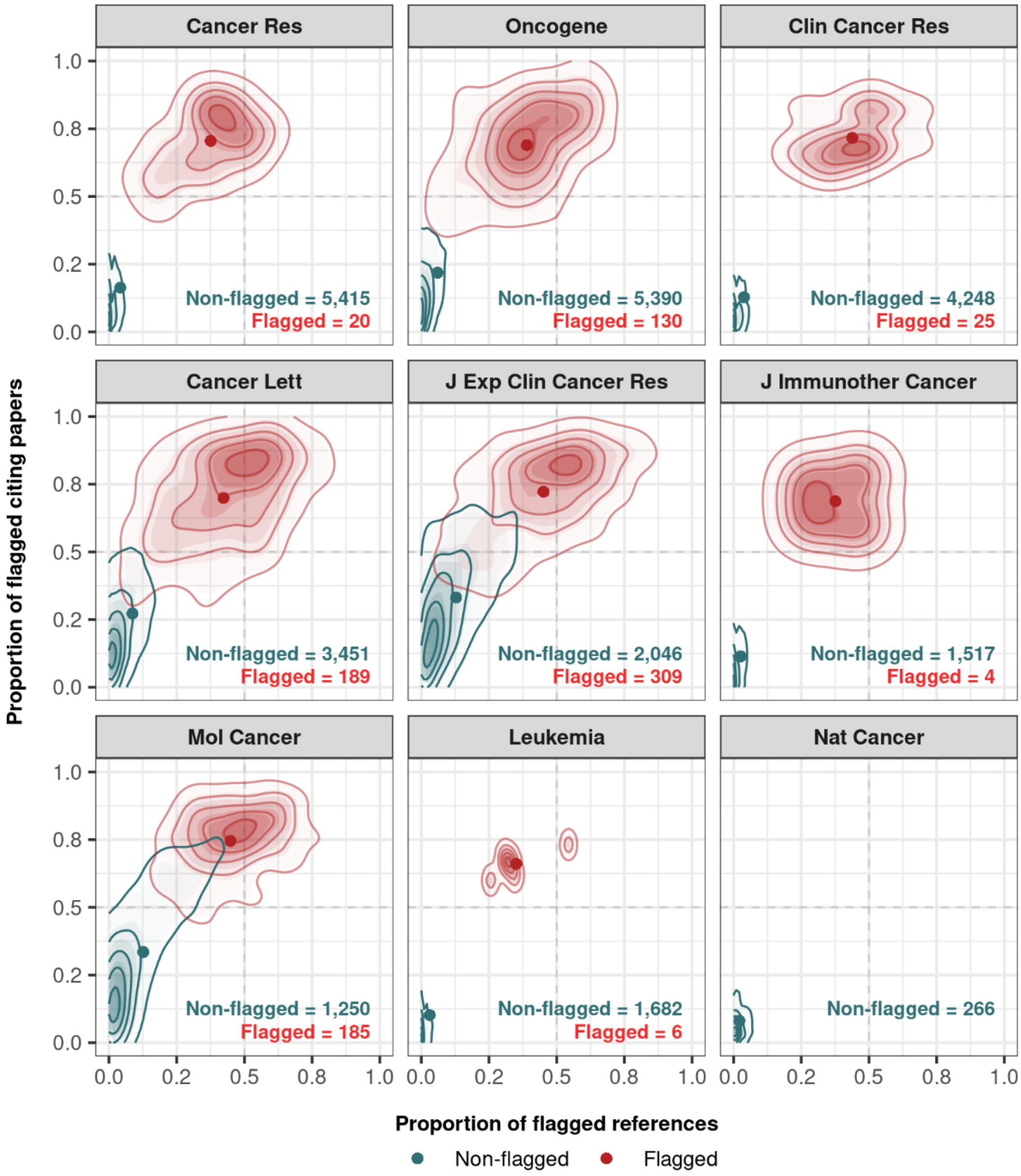


*Fig. S9: Two-dimensional kernel density estimates showing, for each of 9 journals, the joint distribution of the proportion of references that cite flagged papers and the proportion of received citations that originate from flagged papers (both 0.6 paper mill probability threshold), separately for flagged and non-flagged index papers (0.9 threshold). The first eight journals correspond to those with the largest numbers of papers in the dataset. Nature Cancer was included for comparison, as this journal had no flagged papers. Vertical and horizontal dashed lines indicate the midpoints of the X- and Y-axes (0.5). Contour lines show density levels from 10% to 90% of each group's peak density, with points indicating each group's centroid (mean). Journal names follow standard PubMed abbreviations.*

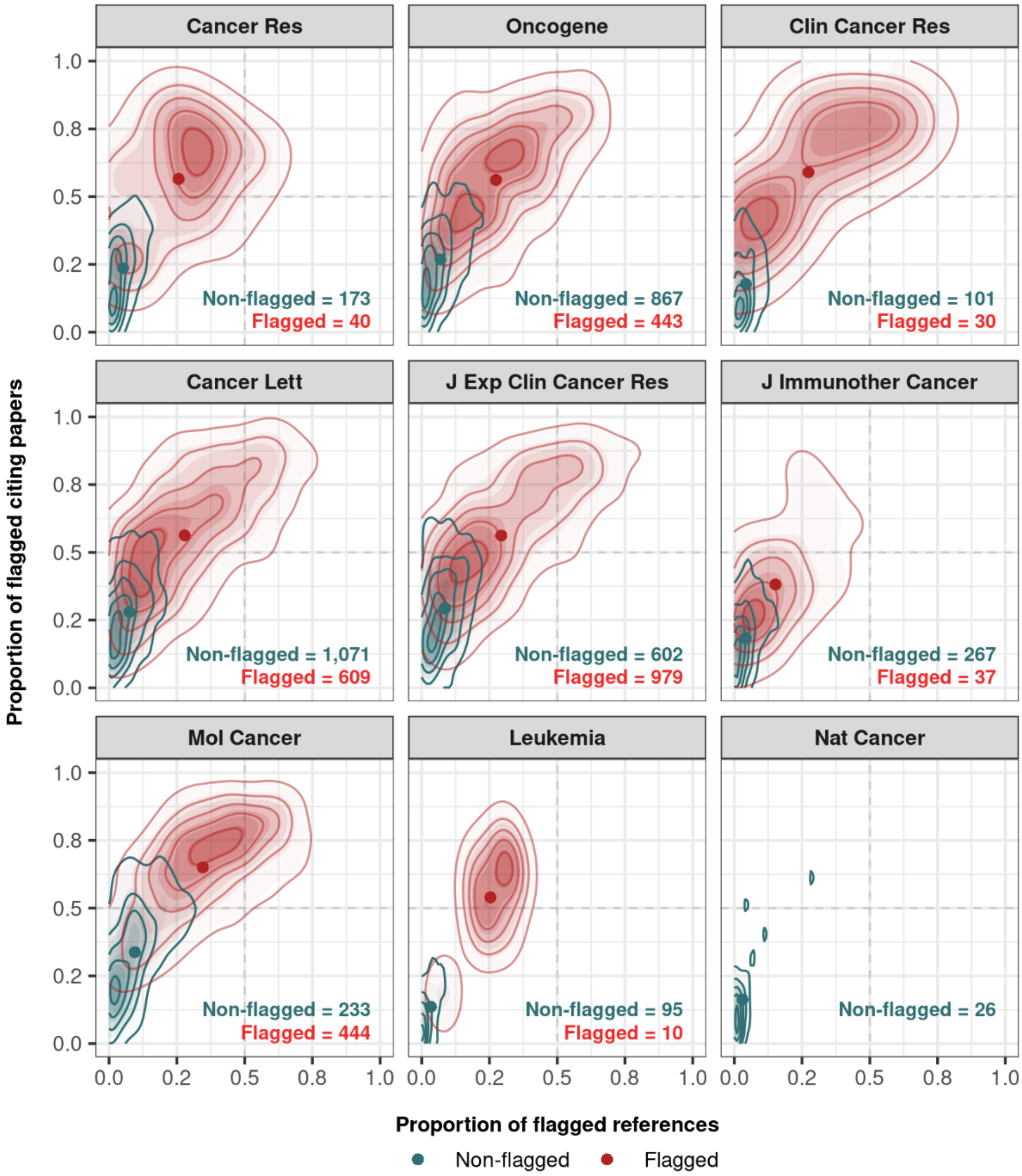


*Fig. S10: Two-dimensional kernel density estimates showing, for China-affiliated index papers in each of 9 journals, the joint distribution of the proportion of references that cite flagged papers and the proportion of received citations that originate from flagged papers, separately for flagged and non-flagged index papers. All results used the 0.6 paper mill probability threshold. The first eight journals correspond to those with the largest numbers of papers in the dataset. Nature Cancer was included for comparison, as this journal had no flagged papers. Vertical and horizontal dashed lines indicate the midpoints of the X- and Y-axes (0.5). Contour lines show density levels from 10% to 90% of each group's peak density, with points indicating each group's centroid (mean). Journal names follow standard PubMed abbreviations.*

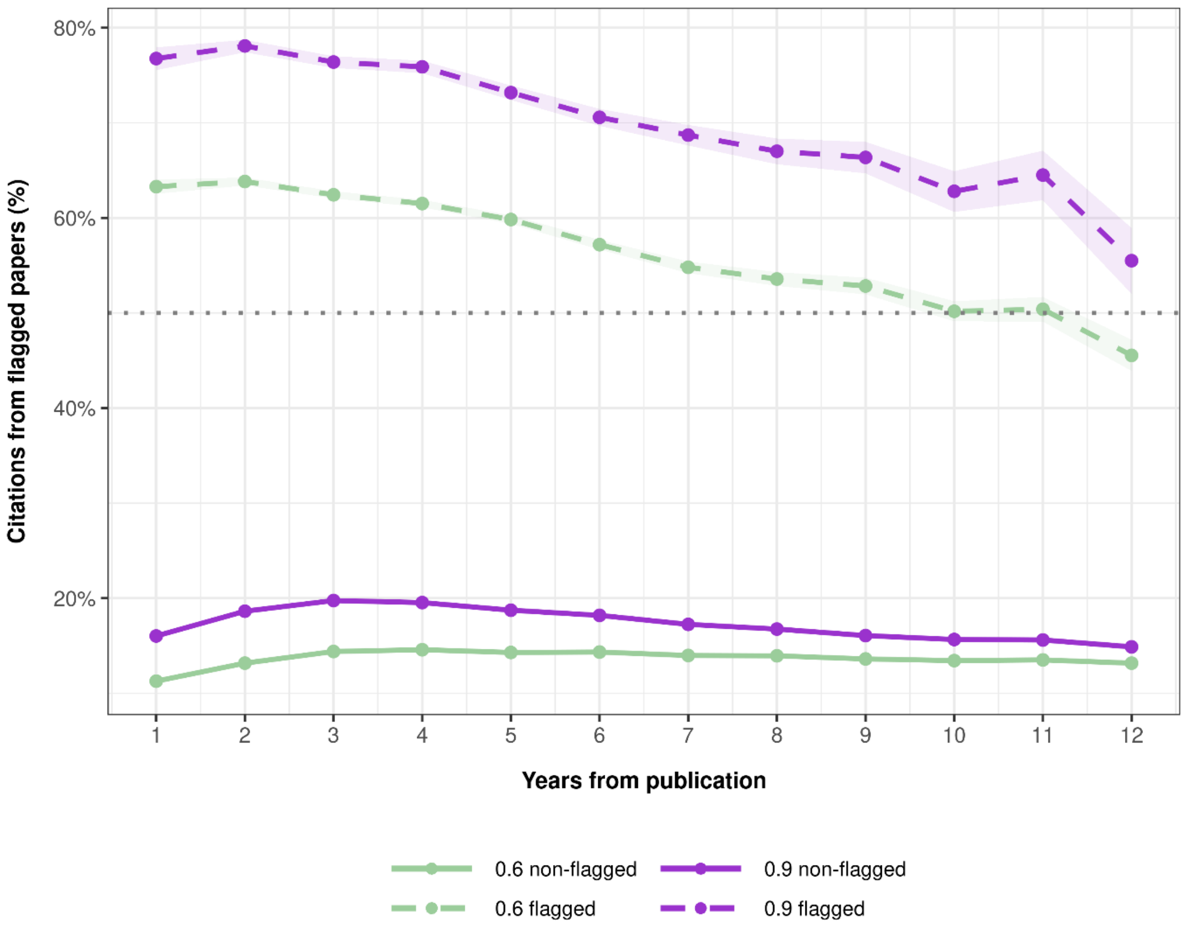


*Fig. S11: Percentages of citations from flagged papers to flagged and non-flagged index papers since publication. Dashed lines represent flagged index papers and solid lines represent non-flagged index papers. Paper mill probability thresholds are represented by green lines (0.6) and purple lines (0.9). The dotted horizontal line indicates 50% of citations from flagged papers. Shaded areas represent 95% confidence intervals.*

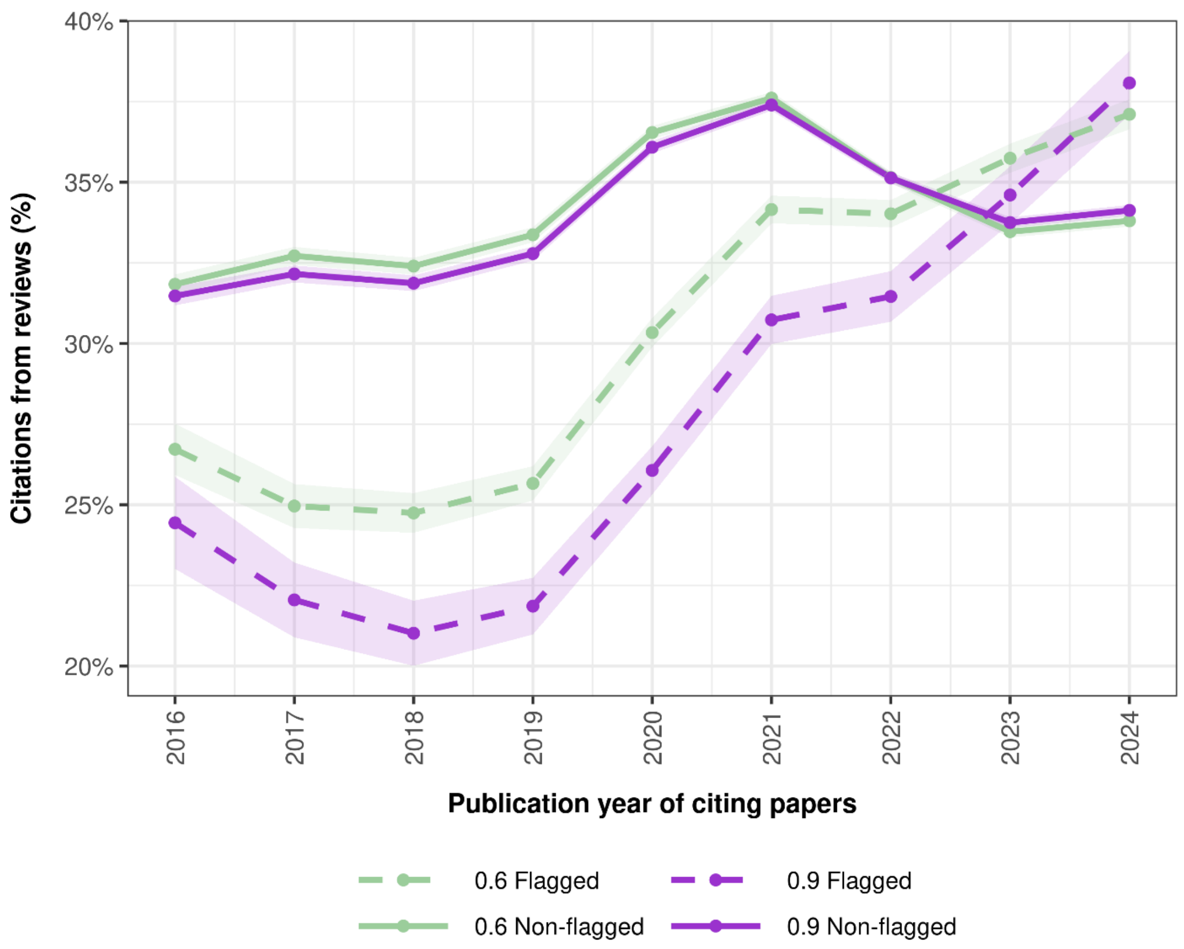


*Fig. S12: Percentages of citations originating from literature reviews to flagged and non-flagged index papers at the 0.6 paper mill probability threshold (green) and the 0.9 threshold (purple). Literature reviews were identified by searching for the "review" keyword in the PubMed publication type metadata. Solid lines represent non-flagged papers whereas dashed lines represent flagged papers. The X-axis is the publication year of citing papers and the Y-axis represents the share of literature reviews among all publication types in each year. The year axis starts at the first year with more than 100,000 citations to ensure stable yearly estimates. 95% confidence intervals are represented by shading.*

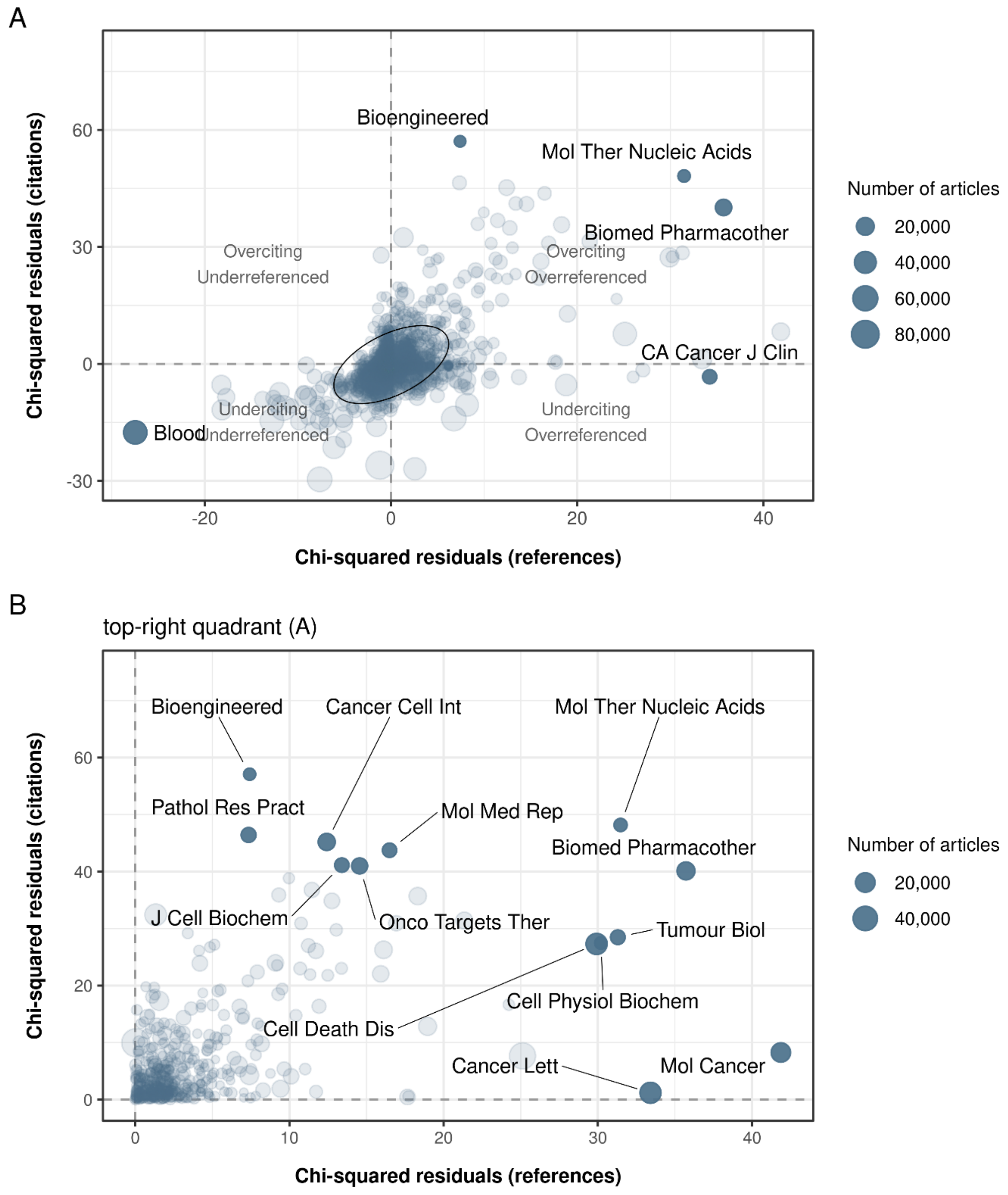


*Fig. S13: Association of journals with citations to (Y-axis) and references from (X-axis) flagged index papers, as determined by Pearson chi-squared residuals at the 0.9 threshold. Each point represents a journal, sized proportionally to the total volume of citations and references linked to flagged index papers. Self-citations and self-references were excluded. Journal names follow standard PubMed abbreviations.* ***(A)*** *All journals positively and / or negatively associated with citations to, and references from, flagged index papers are shown. The central ellipse shows a dense area covering 95% of journals where both residuals are relatively close to zero; grey points represent all remaining journals.* ***(B)*** *Journals showing excess citations to and references from flagged index papers are displayed (**(A)**, top-right quadrant). Labelled points correspond to the ten journals most overrepresented in citations and the ten most overrepresented in references from flagged index papers; some journals are in both groups.*

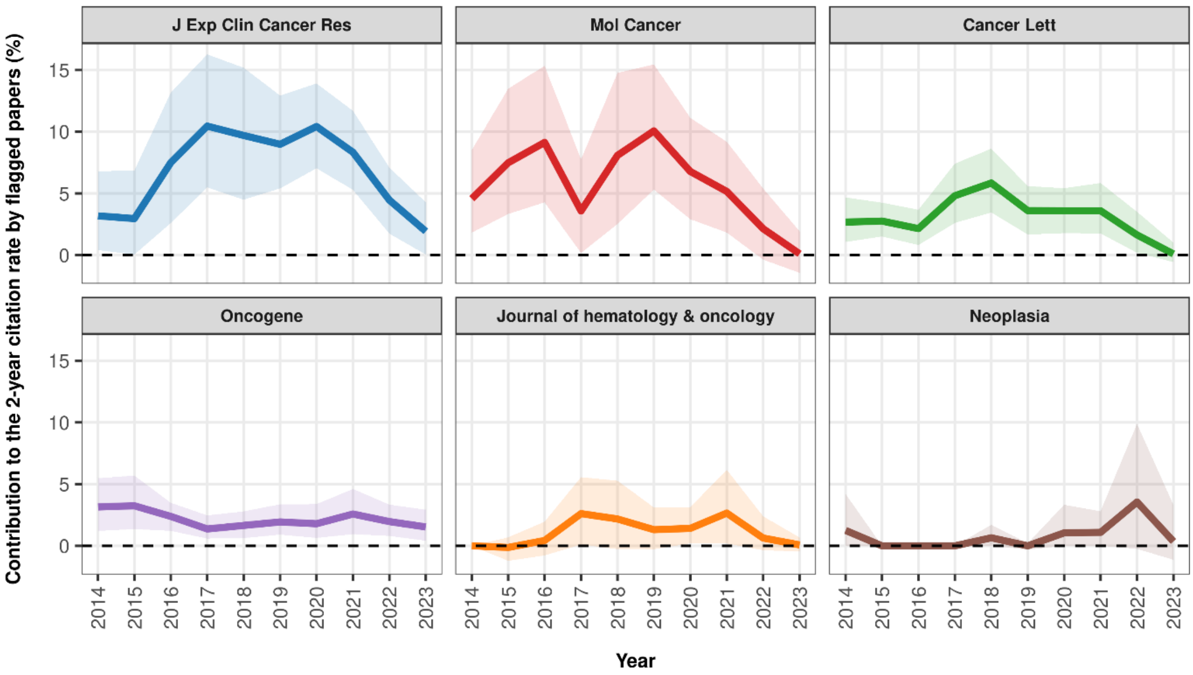


*Fig. S14: Contribution (%) of flagged index papers to the 2-year citation rate of 6 journals (Y-axis) at the 0.9 paper mill probability threshold, for each year of calculation (X-axis). For example, a value of 5% indicates that the journal's 2-year citation rate would have been approximately 5% lower in the absence of flagged papers and their citations. Values below zero would indicate years in which flagged papers contributed less to the 2-year citation rate than non-flagged papers. 95% confidence intervals are shown as shaded bands around the lines.*

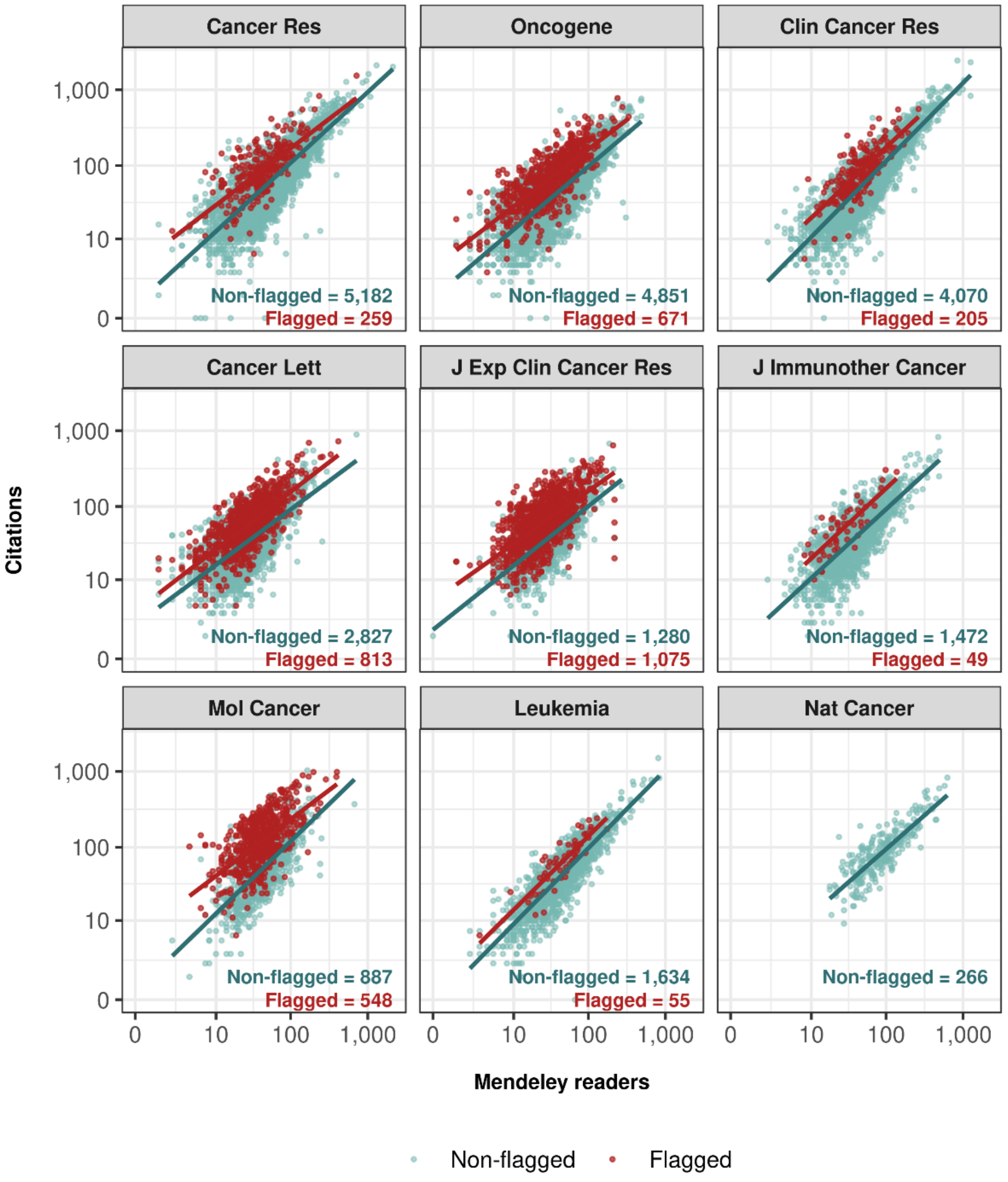


*Fig. S15: Association between citation counts and Mendeley readers across flagged and non-flagged papers in 9 journals at the 0.6 paper mill probability threshold. Both variables are presented on a log10 scale. The first eight journals correspond to those with the largest numbers of papers in the dataset. Nature Cancer was included for comparison, as this journal had no flagged papers. Each point represents an individual paper, and solid lines indicate fitted linear regression models stratified by flagged and non-flagged papers. Journal names follow standard PubMed abbreviations. Same as Fig. S5, showing all values.*

#### Supplementary Tables

*Table S1: Number of molecular papers from leading cancer journals published from 2012 to 2023. Journals are sorted by numbers of selected molecular cancer research papers. The total number of papers published over the same period is also displayed (including non-molecular research papers, reviews, editorials, letters, etc.).*

| Journal | Publisher | Selected papers | Total papers |
|---|---|---|---|
| Oncogene | Springer Nature | 5,522 | 6,160 |
| Cancer Research | American Association for Cancer Research | 5,441 | 6,643 |
| Clinical Cancer Research | American Association for Cancer Research | 4,275 | 7,910 |
| Cancer Letters | Elsevier | 3,640 | 5,188 |
| Journal of Experimental & Clinical Cancer Research | Springer Nature | 2,355 | 2,887 |
| Leukemia | Springer Nature | 1,689 | 2,736 |
| Journal for ImmunoTherapy of Cancer | BMJ Group | 1,521 | 2,773 |
| Molecular Cancer | Springer Nature | 1,435 | 2,004 |
| Cancer Cell | Elsevier | 1,142 | 2,045 |
| Cancer Immunology Research | American Association for Cancer Research | 1,068 | 1,369 |
| Neoplasia | Elsevier | 1,012 | 1,158 |
| Cancer Discovery | American Association for Cancer Research | 985 | 3,578 |
| Neuro-Oncology | Oxford University Press | 798 | 1,875 |
| Journal of Hematology & Oncology | Springer Nature | 634 | 1,621 |
| Journal of Thoracic Oncology | Elsevier | 514 | 2,093 |
| NPJ Breast Cancer | Springer Nature | 276 | 594 |
| NPJ Precision Oncology | Springer Nature | 272 | 449 |
| Nature Cancer | Springer Nature | 266 | 350 |
| Experimental Hematology & Oncology | Springer Nature | 180 | 564 |
| Cancer Communications | John Wiley & Sons | 134 | 342 |

*Table S2: Number and percentage of flagged index papers in the selected journals, based on 0.6 and 0.9 paper mill probability thresholds. Percentages use the flagged papers among the total index papers published in each journal. The journals are ordered by the number of flagged index papers at the 0.6 threshold.*

| Journal | Flagged papers (p = 0.6) | Flagged papers (p = 0.9) |
|---|---|---|
| Journal of Experimental & Clinical Cancer Research | 1,075 (46%) | 309 (13%) |
| Cancer Letters | 813 (22%) | 189 (5%) |
| Oncogene | 671 (12%) | 130 (2%) |
| Molecular Cancer | 548 (38%) | 185 (13%) |
| Cancer Research | 259 (5%) | 20 (<1%) |
| Clinical Cancer Research | 205 (5%) | 25 (1%) |
| Journal of Hematology & Oncology | 117 (19%) | 31 (5%) |
| Neuro-Oncology | 72 (9%) | 15 (2%) |
| Neoplasia | 67 (7%) | 7 (1%) |
| Leukemia | 55 (3%) | 6 (<1%) |
| Cancer Communications | 49 (37%) | 3 (2%) |
| Journal for ImmunoTherapy of Cancer | 49 (3%) | 4 (<1%) |
| Experimental Hematology & Oncology | 33 (18%) | 3 (2%) |
| Journal of Thoracic Oncology | 18 (4%) | 1 (<1%) |
| Cancer Cell | 17 (2%) | 2 (<1%) |
| Cancer Immunology Research | 15 (1%) | 1 (<1%) |
| NPJ Breast Cancer | 9 (3%) | 1 (<1%) |
| NPJ Precision Oncology | 8 (3%) | 1 (<1%) |
| Cancer Discovery | 5 (<1%) | 0 (0%) |
| Nature Cancer | 0 (0%) | 0 (0%) |

Percentages below 1% are reported as <1%.

*Table S3: Number and percentage of flagged index papers by country of affiliation of authors for the ten countries with the highest numbers of flagged index papers, using the 0.6 and 0.9 paper mill probability thresholds. Percentages are for flagged papers among all papers published by each country in the selected journals. Rows are ordered by the number of flagged papers at the 0.6 probability threshold.*

| Country | Flagged papers (p = 0.6) | Flagged papers (p = 0.9) |
|---|---|---|
| China | 2,845 (41%) | 811 (12%) |
| United States | 336 (4%) | 36 (<1%) |
| Japan | 86 (9%) | 7 (<1%) |
| Taiwan | 83 (15%) | 12 (2%) |
| South Korea | 80 (11%) | 9 (1%) |
| Italy | 78 (7%) | 5 (<1%) |
| Germany | 70 (5%) | 4 (<1%) |
| Hong Kong | 25 (29%) | 1 (1%) |
| Spain | 22 (4%) | 0 (0%) |
| Canada | 18 (3%) | 0 (0%) |

Percentages below 1% are reported as <1%.

*Table S4: Number of citations and attention metrics according to the flagged status of index papers for both probability levels. Average numbers of citations received (total, 1 year, and 3 years after publication) are shown along with average Mendeley readers and online accesses. 95% confidence intervals are shown in brackets below each value.*

| prob. | index papers | n | avg cites | avg cites (1 years) | avg cites (3 years) | avg mendeley readers | avg online accesses |
|---|---|---|---|---|---|---|---|
| 0.6 | Non-flagged | 29,074 (87.7%) | 80.7<br>(79.4–82.1) | 3.5<br>(3.4–3.6) | 25<br>(24.6–25.3) | 67<br>(66–68) | 4,112<br>(4,027–4,197) |
| 0.6 | Flagged | 4,085 (12.3%) | 96.7<br>(93.5–99.8) | 4.2<br>(4.0–4.4) | 36.6<br>(35.5–37.7) | 41<br>(40–42) | 5,108<br>(4,973–5,243) |
| 0.9 | Non-flagged | 32,226 (97.3%) | 81.7<br>(80.4–82.9) | 3.5<br>(3.5–3.6) | 25.8<br>(25.5–26.1) | 64<br>(63–65) | 4,206<br>(4,127–4,284) |
| 0.9 | Flagged | 933 (2.7%) | 118.4<br>(110.9–125.9) | 5.6<br>(5.1–6.1) | 46.7<br>(44.0–49.4) | 40<br>(37–42) | 5,178<br>(4,957–5,400) |

Online access data were available for only 14 journals; therefore, this subset does not cover the same population as the other metrics.

*Table S5: Counts and percentages of flagged and non-flagged citing papers, stratified by the predicted status of index papers at two probability thresholds (p = 0.6 and p = 0.9). Index papers were classified using the two probability thresholds, while citing papers were screened using the lower threshold of p = 0.6. Counts and percentages represent the number and proportion of citing papers within each index paper category. Empty abstracts were removed from this visualisation.*

| Probability | Index paper | Citing paper | Count | Percentage |
|---|---|---|---|---|
| 0.6 | Non-flagged | Non-flagged | 1,775,551 | 86.2% |
| | | Flagged | 283,073 | 13.8% |
| | Flagged | Non-flagged | 131,227 | 39.6% |
| | | Flagged | 199,775 | 60.4% |
| 0.9 | Non-flagged | Non-flagged | 1,882,930 | 82% |
| | | Flagged | 413,713 | 18% |
| | Flagged | Non-flagged | 23,848 | 25.6% |
| | | Flagged | 69,135 | 74.4% |

*Table S6: Counts and percentages of flagged and non-flagged referenced papers, stratified by the predicted status of index papers at two probability thresholds (p = 0.6 and p = 0.9). Index papers were classified using the two probability thresholds, while referenced papers were screened using the lower threshold of p = 0.6. Counts and percentages represent the number and proportion of referenced papers within each index paper category. Empty abstracts were removed from this visualisation.*

| Probability | Index paper | Referenced paper | Count | Percentage |
|---|---|---|---|---|
| 0.6 | Non-flagged | Non-flagged | 1,308,627 | 96.4% |
| | | Flagged | 49,214 | 3.6% |
| | Flagged | Non-flagged | 130,610 | 72.2% |
| | | Flagged | 50,368 | 27.8% |
| 0.9 | Non-flagged | Non-flagged | 1,416,645 | 94.5% |
| | | Flagged | 82,721 | 5.5% |
| | Flagged | Non-flagged | 22,592 | 57.3% |
| | | Flagged | 16,861 | 42.7% |

*Table S7: Country of affiliation of papers citing index papers, stratified by index paper status (non-flagged and flagged) and probability thresholds. Countries are ranked in descending order by number of citations. Percentages represent each country's contribution to the total number of citations within a given group (e.g., non-flagged papers with a paper mill probability threshold of 0.6).*

| Probability | Index paper | Origin of citations | Citations | Percentage |
|---|---|---|---|---|
| 0.6 | Non-flagged | China | 673,615 | 30.7% |
| | | United States | 551,617 | 25.1% |
| | | Italy | 93,783 | 4.3% |
| | | Germany | 84,300 | 3.8% |
| | | United Kingdom | 70,360 | 3.2% |
| | Flagged | China | 218,642 | 62.5% |
| | | United States | 28,715 | 8.2% |
| | | Iran, Islamic Republic of | 12,291 | 3.5% |
| | | India | 10,521 | 3% |
| | | Italy | 8,961 | 2.6% |
| 0.9 | Non-flagged | China | 825,928 | 33.7% |
| | | United States | 574,582 | 23.5% |
| | | Italy | 100,855 | 4.1% |
| | | Germany | 88,625 | 3.6% |
| | | United Kingdom | 73,123 | 3% |
| | Flagged | China | 66,329 | 68.7% |
| | | United States | 5,750 | 6% |
| | | Iran, Islamic Republic of | 4,062 | 4.2% |
| | | India | 2,511 | 2.6% |
| | | Italy | 1,889 | 2% |

*Table S8: Country of affiliation of papers referenced by index papers, stratified by index paper status (non-flagged and flagged) and probability thresholds. Countries are ranked in descending order by number of references. Percentages represent each country's contribution to the total number of references within a given group (e.g., non-flagged papers with a paper mill probability threshold of 0.6).*

| Probability | Index paper | Country of referenced paper | References | Percentage |
|---:|---|---|---:|---:|
| 0.6 | Non-flagged | United States | 677,878 | 50.4% |
| | | United Kingdom | 84,243 | 6.3% |
| | | China | 77,599 | 5.8% |
| | | Germany | 71,519 | 5.3% |
| | | Japan | 56,372 | 4.2% |
| | Flagged | United States | 67,231 | 37.4% |
| | | China | 43,515 | 24.2% |
| | | Japan | 8,261 | 4.6% |
| | | Germany | 7,805 | 4.3% |
| | | United Kingdom | 6,564 | 3.7% |
| 0.9 | Non-flagged | United States | 731,923 | 49.2% |
| | | China | 108,117 | 7.3% |
| | | United Kingdom | 89,762 | 6% |
| | | Germany | 77,840 | 5.2% |
| | | Japan | 63,059 | 4.2% |
| | Flagged | United States | 13,186 | 33.6% |
| | | China | 12,997 | 33.2% |
| | | Japan | 1,574 | 4% |
| | | Germany | 1,484 | 3.8% |
| | | United Kingdom | 1,045 | 2.7% |

*Table S9: Journals overrepresented in citations to and references from flagged index papers, which are also associated with known integrity and paper mill concerns.*

| **Journal** | **Known concern(s)** |
|---|---|
| Aging | Listed in Beall's list (9); High number of suspected paper mill products (10) |
| Molecular Therapy – Nucleic Acids | Paper mill target (11); Retraction campaign (12) |
| Bioengineered | Delisted from Clarivate (13); Paper mill target (14–16) |
| Cancer Cell International | Paper mill target (17); Paper mill-associated retractions (18) |
| Pathology – Research and Practice | Fake peer review (19) |
| Tumor Biology | Fake peer review (20) |
| Cell Death & Disease | Image manipulation (21) |
| Molecular Medicine Reports | Paper mill-associated retractions (22); High number of suspected paper mill products (10) |
| Biomedicine & Pharmacotherapy | Paper mill-associated retractions (22); High number of suspected paper mill products (10) |
| Journal of Cellular Biochemistry | Paper mill-associated retractions (23); Paper mill target (24); High number of suspected paper mill products (10) |
| Molecular Cancer | Wrongly identified nucleotide sequences (25) |
| Oncology Reports | High number of suspected paper mill products (10) |
| OncoTargets and Therapy | High number of suspected paper mill products (10) |

*Table S10: Number of citations received by all flagged index papers for both probability thresholds. The table rows are ordered according to the total number of citations received by flagged index papers at the p = 0.6 threshold. Citation counts were collected up to February 2026. Zero values indicate that either no index papers were flagged at the corresponding threshold or that flagged index papers received no citations. Percentages relative to the total number of citations are shown in brackets.*

| Journal | Citations to flagged papers (p=0.6) | Citations to flagged papers (p=0.9) |
|---|---|---|
| Molecular Cancer | 86,565 (57%) | 33,900 (22%) |
| Journal of Experimental & Clinical Cancer Research | 84,497 (57%) | 26,398 (18%) |
| Cancer Letters | 62,042 (32%) | 17,838 (9%) |
| Oncogene | 60,484 (17%) | 14,523 (4%) |
| Cancer Research | 30,717 (6%) | 3,624 (1%) |
| Clinical Cancer Research | 21,872 (6%) | 3,056 (1%) |
| Journal of Hematology & Oncology | 14,703 (33%) | 4,601 (10%) |
| Cancer Cell | 7,140 (2%) | 2,727 (1%) |
| Neuro-Oncology | 5,782 (11%) | 1,742 (3%) |
| Leukemia | 4,510 (4%) | 771 (1%) |
| Neoplasia | 4,105 (10%) | 491 (1%) |
| Journal for ImmunoTherapy of Cancer | 3,313 (4%) | 254 (<1%) |
| Cancer Communications | 3,094 (44%) | 196 (3%) |
| Cancer Immunology Research | 1,492 (2%) | 50 (<1%) |
| Journal of Thoracic Oncology | 1,422 (4%) | 127 (<1%) |
| Experimental Hematology & Oncology | 1,343 (28%) | 138 (3%) |
| Cancer Discovery | 1,156 (1%) | 0 (0%) |
| NPJ Breast Cancer | 361 (4%) | 30 (<1%) |
| NPJ Precision Oncology | 258 (3%) | 12 (<1%) |
| Nature Cancer | 0 (0%) | 0 (0%) |

*Table S11: Location and specification of all statistical models.*

| Location | Test | Dataset | Outcome | Model specification | SE estimation |
|---|---|---|---|---|---|
| Fig. 3 | Negative Binomial Regression | Index Papers | Total Citations | TC = IPS + Year + Journal | Cluster-robust (Journal×Year) |
| Fig. 3 | Negative Binomial Regression | Index Papers | Early Citations (1 year) | EC1 = IPS + Year + Journal | Cluster-robust (Journal×Year) |
| Fig. 3 | Negative Binomial Regression | Index Papers | Early Citations (3 years) | EC3 = IPS + Year + Journal | Cluster-robust (Journal×Year) |
| Fig. 3 | Negative Binomial Regression | Index Papers | Mendeley Readers | MR = IPS + Year + Journal | Cluster-robust (Journal×Year) |
| Fig. 3 | Negative Binomial Regression | Index Papers | Online Accesses | OA = IPS + Year + Journal | Cluster-robust (Journal×Year) |
| Fig. S2 | Negative Binomial Regression | Index Papers | Total Citations | TC = IPS + Year + Journal + Country | Cluster-robust (Journal×Year) |
| Fig. S2 | Negative Binomial Regression | Index Papers | Early Citations (1 year) | EC1 = IPS + Year + Journal + Country | Cluster-robust (Journal×Year) |
| Fig. S2 | Negative Binomial Regression | Index Papers | Early Citations (3 years) | EC3 = IPS + Year + Journal + Country | Cluster-robust (Journal×Year) |
| Fig. S2 | Negative Binomial Regression | Index Papers | Mendeley Readers | MR = IPS + Year + Journal + Country | Cluster-robust (Journal×Year) |
| Fig. S2 | Negative Binomial Regression | Index Papers | Online Accesses | OA = IPS + Year + Journal + Country | Cluster-robust (Journal×Year) |
| Fig. S3 | Negative Binomial Regression | Dimensions.ai data | Total Citations | TC = IPS + Year + Journal | Cluster-robust (Journal×Year) |
| Fig. S3 | Negative Binomial Regression | Dimensions.ai data | Early Citations (1 year) | EC1 = IPS + Year + Journal | Cluster-robust (Journal×Year) |
| Fig. S3 | Negative Binomial Regression | Dimensions.ai data | Early Citations (3 years) | EC3 = IPS + Year + Journal | Cluster-robust (Journal×Year) |
| Fig. S3 | Negative Binomial Regression | Dimensions.ai data | Mendeley Readers | MR = IPS + Year + Journal | Cluster-robust (Journal×Year) |
| Fig. S3 | Negative Binomial Regression | Dimensions.ai data | Online Accesses | OA = IPS + Year + Journal | Cluster-robust (Journal×Year) |
| Fig. S4 (A & B) | Negative Binomial Regression | ncRNA Index Papers | Total Citations | TC = IPS + Year + Journal ± Country | Cluster-robust (Journal×Year) |
| Fig. S4 (A & B) | Negative Binomial Regression | ncRNA Index Papers | Early Citations (1 year) | EC1 = IPS + Year + Journal ± Country | Cluster-robust (Journal×Year) |
| Fig. S4 (A & B) | Negative Binomial Regression | ncRNA Index Papers | Early Citations (3 years) | EC3 = IPS + Year + Journal ± Country | Cluster-robust (Journal×Year) |
| Fig. S4 (A & B) | Negative Binomial Regression | ncRNA Index Papers | Mendeley Readers | MR = IPS + Year + Journal ± Country | Cluster-robust (Journal×Year) |
| Fig. S4 (A & B) | Negative Binomial Regression | ncRNA Index Papers | Online Accesses | OA = IPS + Year + Journal ± Country | Cluster-robust (Journal×Year) |
| Results, citation clustering | Mixed-Effects Logistic Regression | Citing Papers | Citing Paper Status | CPS = IPS + Year + Journal + (1 \| IPD) | Default |
| Results, citation clustering | Mixed-Effects Logistic Regression | Referenced Papers | Referenced Paper Status | RPS = IPS + Year + Journal + (1 \| IPD) | Default |
| Results, citation clustering (Country adjustment) | Mixed-Effects Logistic Regression | Citing Papers | Citing Paper Status | CPS = IPS + Year + Journal + Country + (1 \| IPD) | Default |
| Results, citation clustering (Country adjustment) | Mixed-Effects Logistic Regression | Referenced Papers | Referenced Paper Status | RPS = IPS + Year + Journal + Country + (1 \| IPD) | Default |
| Results, citation clustering (out-of-domain sensitivity analysis) | Mixed-Effects Logistic Regression | Citing Papers from 20 Index Journals | Citing Paper Status | CPS = IPS + Year + Journal + (1 \| IPD) | Default |
| Results, citation clustering (out-of-domain sensitivity analysis) | Mixed-Effects Logistic Regression | Referenced Papers from 20 Index Journals | Referenced Paper Status | RPS = IPS + Year + Journal + (1 \| IPD) | Default |
| Results, citation clustering (topical homophily sensitivity analysis) | Mixed-Effects Logistic Regression | Citing Papers | Citing Paper Status | CPS = IPS + Year + Journal + KCC + (1 \| IPD) | Default |
| Results, citation clustering (topical homophily sensitivity analysis) | Mixed-Effects Logistic Regression | Referenced Papers | Referenced Paper Status | RPS = IPS + Year + Journal + KCC + (1 \| IPD) | Default |
| Fig. 5 | Pearson Chi-Squared Test | Citing Papers | Journal Overrepresentation | Citation Counts × Journal × IPS | - |
| Fig. 5 | Pearson Chi-Squared Test | Referenced Papers | Journal Overrepresentation | Citation Counts × Journal × IPS | - |
| Fig. S13 | Pearson Chi-Squared Test | Citing Papers | Journal Overrepresentation | Citation Counts × Journal × IPS | - |
| Fig. S13 | Pearson Chi-Squared Test | Referenced Papers | Journal Overrepresentation | Citation Counts × Journal × IPS | - |

Paper Status refers to the model classification of a paper (0 = Non-Flagged / 1 = Flagged). Country is a binary indicator of authorship origin (0 = non-Chinese / 1 = Chinese). IPS = Index Paper Status, CPS = Citing Paper Status, RPS = Referenced Paper Status, IPD = Index Paper DOI, and KCC = K-Cluster Classification (with K in range [5, 10, 20, 30, 50, 100]). (1 | IPD) indicates a random intercept.

#### Supplementary Datasets

**Dataset S1:** Molecular oncology journal selection within the top 10% of SCImago's pooled "cancer research" and "oncology" rankings – rankings based on the citations per document (2 years) metric. Molecular oncology journals are highlighted in green, non-molecular oncology journals in red.

**Dataset S2:** Two-year citation rate (2YCR) data for all 20 journals. The sheet includes the full 2YCR data, the 2YCR excluding 0.6-flagged papers, the 2YCR excluding 0.9-flagged papers, and their confidence intervals. The "SCImago Impact Factor" corresponds to the citations per document (2 years) metric.
